# Connectome-based predictions reveal developmental change in the functional architecture of sustained attention and working memory

**DOI:** 10.1101/2021.08.01.454530

**Authors:** Omid Kardan, Andrew J. Stier, Carlos Cardenas-Iniguez, Julia C. Pruin, Kathryn E. Schertz, Yuting Deng, Taylor Chamberlain, Wesley J. Meredith, Xihan Zhang, Jillian E. Bowman, Tanvi Lakhtakia, Lucy Tindel, Emily W. Avery, Qi Lin, Kwangsun Yoo, Marvin M. Chun, Marc G. Berman, Monica D. Rosenberg

**Affiliations:** University of Chicago; University of California, Los Angeles; Yale University

**Keywords:** sustained attention, working memory, fMRI, functional connectivity, predictive modeling, child development

## Abstract

Sustained attention (SA) and working memory (WM) are critical processes, but the brain networks supporting these abilities in development are unknown. We characterized the functional brain architecture of SA and WM in 9–11-year-old children and adults. First, we found that adult network predictors of SA generalized to predict individual differences and fluctuations in SA in youth. A WM network model predicted WM performance both across and within children—and captured individual differences in later recognition memory—but underperformed in youth relative to adults. We next characterized functional connections differentially related to SA and WM in youth compared to adults. Results revealed two network configurations: a dominant architecture predicting performance in both age groups and a secondary architecture, more prominent for WM than SA, predicting performance in one. Thus, functional connectivity predicts SA and WM in youth, with networks predicting WM changing more from preadolescence to adulthood than those predicting SA.

## Introduction

Maintaining focus over time and information in working memory—related but separable functions (Gazzaley, & Nobre, 2012; Hollingworth, & Maxcey-Richard, 2013; Xu, et al., 2014; Brissenden, & Somers, 2019; LaBar et al., 1999)—are foundational cognitive processes critical for successfully performing everyday activities across the lifespan. In addition to being integral to everyday life, these cognitive processes vary greatly across individuals (Rosenberg et al., 2017; Fortenbaugh et al., 2015; Luck et al., 2013) and fluctuate over time within the same person (Adam & deBettencourt, 2019). These inter- and intra-individual differences are particularly important to study in development because of their consequences for life-long achievements. For example, research in children and adolescents has suggested that attention is more predictive of later academic achievement than more general problem behaviors (e.g., aggression and non-compliance) and interpersonal skills (Barriga et al., 2002; Hinshaw, 1992; Rabiner, et al., 2016).

Network neuroscience proposes that cognitive and attentional processes are emergent properties of interactions between brain regions (Bassett & Sporns, 2017). The success of recent work predicting behavior based on functional magnetic resonance imaging (fMRI) functional connectivity (i.e., the correlation between synchronous BOLD activity among pairs of brain regions) supports the tenability of this position (Yamashita et al., 2018; Smith et al., 2015; Rosenberg et al., 2016a, Rosenberg et al., 2020; Song et al., 2021). In other words, this work suggests that the degree to which activity is coordinated across large-scale brain networks may better characterize cognitive processes than the magnitude of activity in single regions in isolation (Smith et al., 2013).

Therefore, in the current study we aimed to understand the development of sustained attention (SA) and working memory (WM) through the lens of network neuroscience. To do this, we used two approaches. First, we assessed the degree to which connectome-based predictive models of SA and WM defined in adults (i.e., mature networks) generalize to predict SA and WM in preadolescents. Second, we characterized the functional brain connections that are *differentially* related to SA and WM performance in preadolescents compared to adults, both within the constraints of the adult networks and in a whole-brain data-driven manner.

### Mature connectome-based approach

Despite the popularity of connectome-based predictive modeling of behavior, cross-dataset and cross-population testing is rare. In other words, brain-based predictive models defined in one dataset are rarely validated in other samples, and even less so in other participant populations (e.g., different ages or diagnoses, see Woo et al., 2017). Hence, existing “publication preregistered” brain markers are currently underutilized and underscrutinized, which obscures both their potential and limitations (Marek et al, 2022). Testing the generalizability of connectivity-based models across ages can inform the degree to which adults and children share common network predictors of cognition and delineate models’ predictive boundaries. Cross-age model validation may also provide insight into how networks underlying cognitive and attentional processes change with development. Additionally, validating models of different cognitive processes (e.g., sustained attention and working memory) to evaluate their unique contributions to predicting behavior can determine if distinctions between the models are behaviorally relevant, and whether those distinctions generalize to different stages of development.

To address these gaps, in Study 1 we utilized previously developed neuromarkers in the form of large-scale functional networks defined to predict sustained attention (Rosenberg et al., 2016b) and working memory (Avery et al., 2020) in adults. We applied these adult connectome-based models to data from the Adolescent Brain Cognitive Development^SM^ (ABCD) Study to predict individual differences and block-to-block changes in sustained attention and working memory task performance in youth. In addition, to characterize relationships between sustained attention, working memory, and long-term memory, we asked whether these same models not only predicted ongoing task performance, but also predict subsequent recognition memory for task stimuli. Successful model generalization would suggest that the functional networks underlying sustained attention and working memory overlap between children and adults. Furthermore, a dissociation such that the neuromarker of sustained attention captures sustained attention performance whereas the neuromarker of working memory captures working memory performance would provide evidence that separable networks support these processes in development.

### Direct comparison approach

In the second approach we aimed to characterize the functional connections that were differentially related to sustained attention and working memory performance in children compared to adults. This was performed both within the constraints of the adult networks and in a whole-brain data-driven manner. This approach can complement both Study 1 and also the existing work that has revealed, for example, changes in the coupling of structural and functional connectivity profiles that may support improvements in working memory and executive abilities in adolescence (Baum et al., 2020).

To achieve this, in Study 2, we combined the behavioral and fMRI data from the youth sample ABCD Study^®^ (9–11 years old) and a large adult sample from the Human Connectome Project (HCP; 21–36+ years old). We first investigated the developmental differences in SA within the constraints of adult networks by a) benchmarking the adult models’ fit to novel adults compared to the preadolescents and b) computationally lesioning different components of the adult networks and comparing how much different regions of the networks were contributing to SA in the youths versus novel adults. Second, we used a multivariate method to find the set of connections that differentiated the youth and adult connectomes with regards to SA and WM performances.

## Results

**Study 1 overview**. In Study 1 we tested the generalizability of network models previously defined to predict sustained attention and working memory in adults to children.

In **Study 1.1**, we asked whether the degree to which children expressed functional connectivity markers of sustained attention (Rosenberg et al., 2016b) and working memory (Avery et al., 2020) previously defined in adult data during an in-scanner *n*-back task predicted their task performance (Figure 1). We hypothesized that the sustained attention connectome-based predictive model would predict 0-back task performance because this low-working-memory-load task is essentially a target detection task similar to a continuous performance task (CPT) traditionally used to assess sustained attention (e.g., Robertson et al., 1997). The sustained attention network may or may not predict 2-back task performance: Although working memory and attention fluctuate in tandem in adults (deBettencourt et al., 2018), sustained attention is not sufficient for successful 2-back task performance. We hypothesized that the working memory connectome-based predictive model, on the other hand, would predict 2-back performance, and that model predictions would be more closely related to 2-back than to 0-back performance because successful 2-back (but not 0-back) performance requires the continuous maintenance and updating of items in working memory.

**Figure 1.**
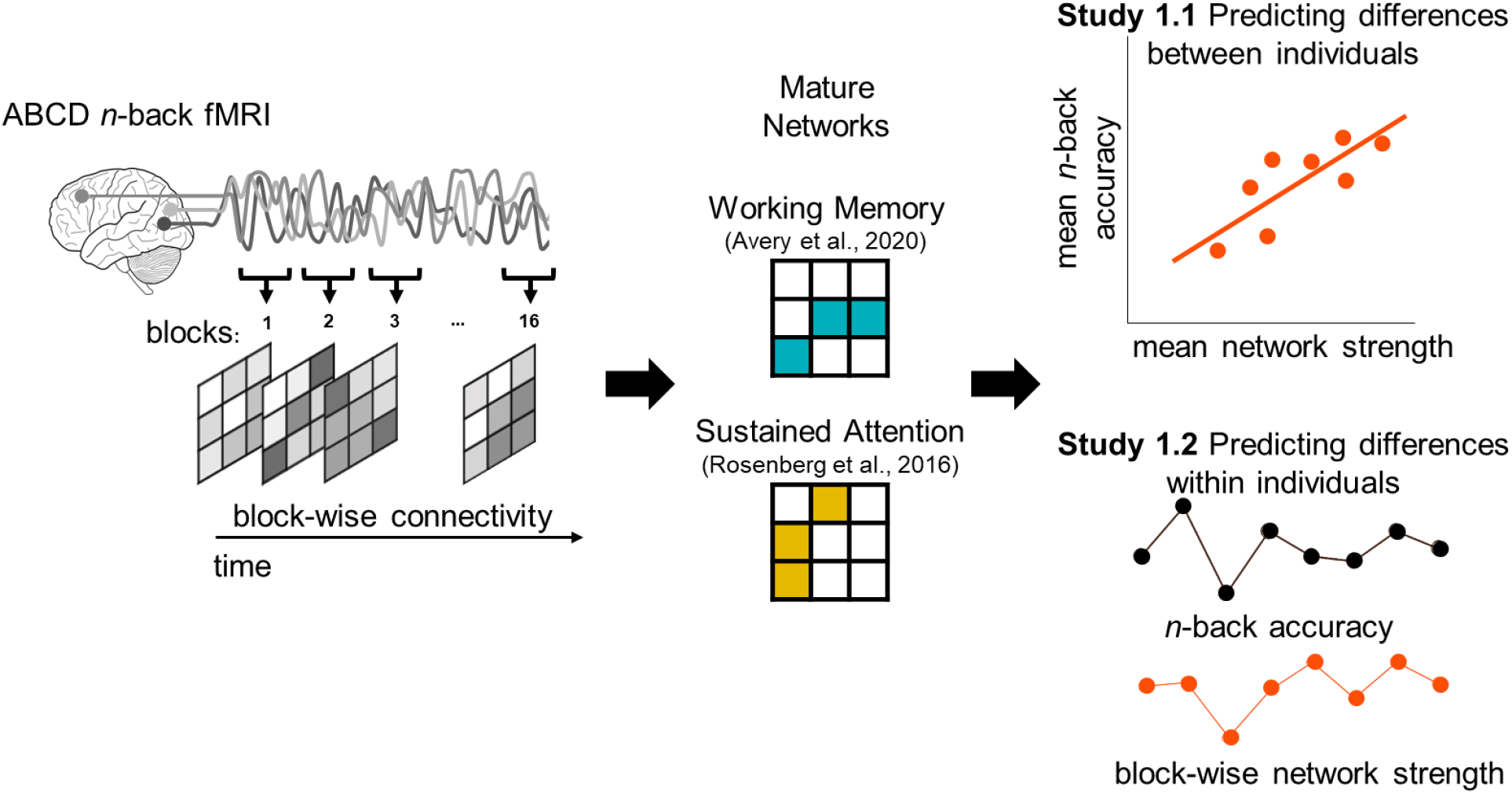
Overview of Study 1. First, we constructed block-wise functional connectivity by correlating blood-oxygen-level-dependent (BOLD) signal time-series from all pairs of functional parcels (left). For each participant, we calculated whole-brain functional connectivity (FC) patterns from fMRI data collected during the eight 0-back and eight 2-back tasks blocks. That is, we calculated up to 16 FC matrices per individual: one using data from each 25-sec (30-31 volumes) *n*-back block separately. Each of the two predefined predictive network masks were then applied to each of these matrices to generate block-specific working memory or sustained attention network strength scores (middle). Each child’s mean network strength during 0-back and 2-back blocks was compared to their mean accuracy in 0-back and 2-back blocks (Study 1.1) or their mean out-of-scanner recognition memory for *n*-back stimuli (Study 1.3). In Study 1.2, block-to-block changes in network strength were compared to corresponding block-to-block changes in 0-back and 2-back accuracy within-subjects.

In **Study 1.2**, we asked whether these same adult network models captured changes in sustained attention and working memory over time in children—that is, whether block-to-block changes in network strength predicted block-to-block changes in *n*-back task accuracy (Figure 1, right panel). Again, we tested for specificity, asking whether the sustained attention and working memory networks better predicted sustained attention (0-back) and working memory (2-back) performance fluctuations, respectively.

Finally, in **Study 1.3**, we asked: Does the degree to which an individual shows a functional connectivity signature of better sustained attention or working memory only affect concurrent task performance, or does it also impact later cognitive processes, such as long-term memory? To investigate this question, we evaluated the consequences of sustained attention and working memory network expression for long-term memory by testing whether network strength during the *n*-back task predicted post-scan recognition memory for task stimuli.

**Study 1.1 Predicting sustained attention and working memory across participants.** Do functional network models defined to predict sustained attention and working memory in adulthood generalize to a large, heterogeneous developmental sample to predict individual differences in these abilities? To test this possibility, we applied our adult connectome-based models of sustained attention and working memory to functional connectivity observed during 9-11-year-olds’ performance of two *n*-back task conditions.

### 1.1.1 Relationship between sustained attention and working memory networks

We hypothesized that the predefined sustained attention (Rosenberg et al., 2016) and working memory (Avery et al., 2020) network masks (Figure 2) capture related but distinct aspects of cognitive function (see *Methods* for descriptions of these networks). Prior to predicting behavioral performance in the ABCD sample, we assessed this hypothesis by *(1)* comparing the anatomy of the sustained attention and working memory network masks, and *(2)* comparing the strength of the sustained attention and working memory networks across participants in the ABCD sample.

**Figure 2.**
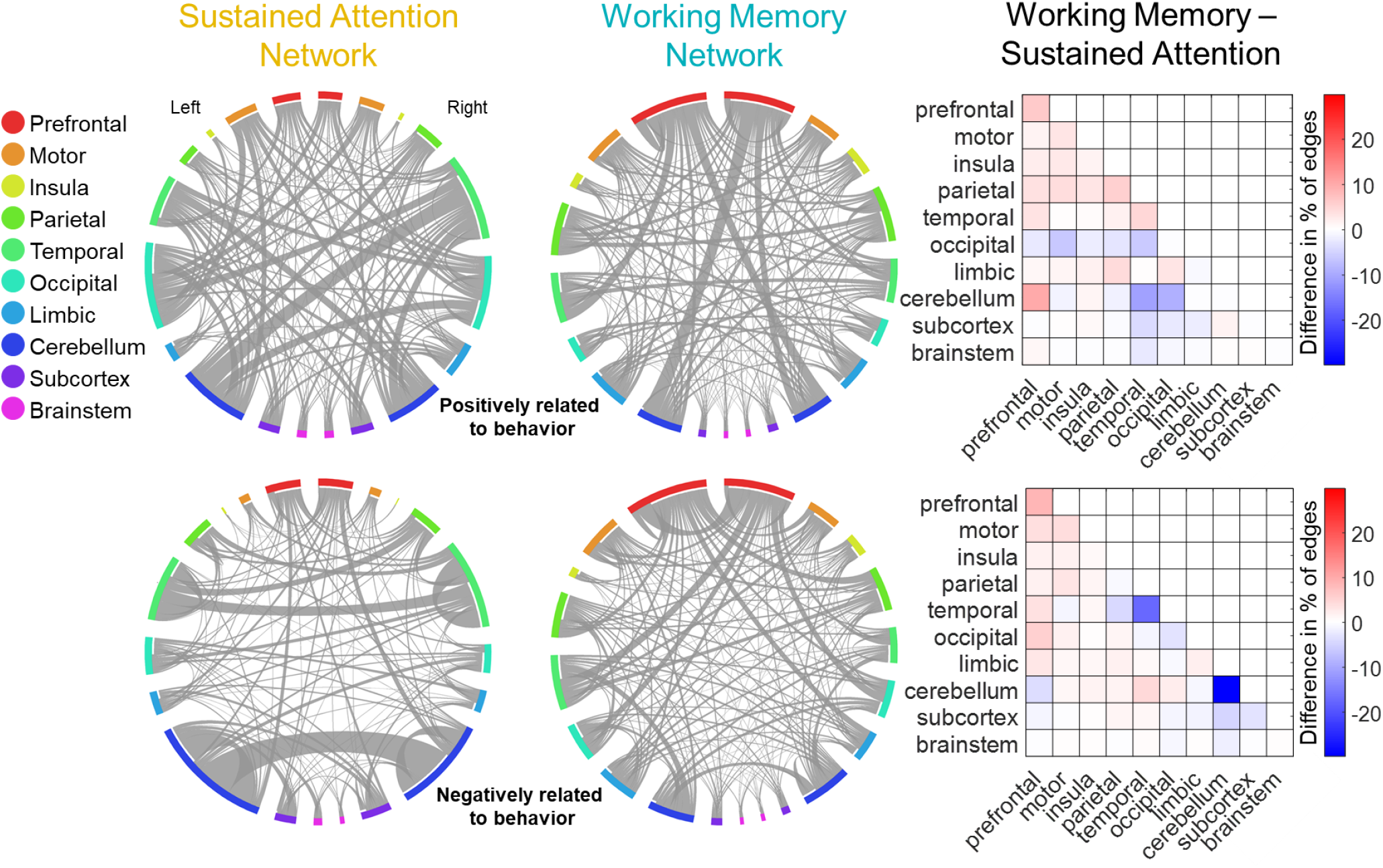
Adult sustained attention and working memory networks and their differences. The circle plots (connectograms) show the sustained attention and working memory networks on the Shen-268 parcels grouped into 20 anatomical regions (10 per hemisphere). The networks positively related to the behavior are shown on the top row and the networks negatively related to the behavior are shown in the bottom row. The matrix plots show the percentage of edges belonging to each macroscale region in working memory connectogram minus the percentage of edges belonging to each macroscale region in sustained attention connectogram. Differences in networks predicting better attention and working memory are shown in the top matrix plot; differences in networks predicting worse attention and working memory are shown in the bottom matrix plot.

First, we found that although the sustained attention and working memory networks both include functional connections, or edges, representing coordinated activity across distributed brain regions, they show little overlap. 37 edges are common to the high-attention and high-working-memory networks (1.5% of combined edges in high-attention and high-working-memory networks, hypergeometric *p* = .351; see *Methods*), which predict better sustained attention and working memory performance, respectively. 33 edges are common to the low-attention and low-working-memory networks (1.8% of combined edges, hypergeometric *p* = .005), which predict worse sustained attention and working memory performance, respectively. Most of these common edges involved prefrontal (32%), motor (21%), and temporal (16%) regions in the high-attention and high-working-memory networks; and cerebellar (45%), occipital (18%), and parietal (18%) regions in the low-attention and low-working-memory networks. There is no significant overlap between the high-attention and low-working-memory networks (19 edges, .9%, *p* = .89) or the low-attention and high-working-memory networks (12 edges, .5%, *p* = .99). At the macroscale region level, the sustained attention networks are more dominated by cerebellar, temporal, and occipital connections whereas the working memory networks include more prefrontal connections (Figure 2).

Anatomical differences between the sustained attention and working memory networks, however, do not guarantee that their strength does not covary together across participants. That is, the degree to which an individual expresses the networks may not be independent. As such, we correlated sustained attention and working memory network strength during the 0-back and 2-back tasks in the ABCD sample (see *Methods*). Briefly, in the 0-back task, children were instructed to detect a target image, shown in the beginning of the block, among a series of images by pressing index versus middle finger on the response box. In the 2-back task, children saw a series of images and determined if the image in each trial matched that of two trials prior to it or not, again by pressing middle versus index finger. In each block, images from one of four categories: faces with positive, negative, and neutral expressions and scenes were used in the task.

Results revealed that sustained attention and working memory network strength values were positively correlated across children during the 0-back task (*r* = .16, *p_adj_* < .001), but negatively correlated during the 2-back task (*r* = –.11, *p_adj_* < .001; black scatterplots in Figure 3). Taken together, the anatomical overlap and network strength correlation analyses suggest that the sustained attention and working memory masks are separable functional networks in children and thus likely do not reflect a monolithic cognitive process.

**Figure 3.**
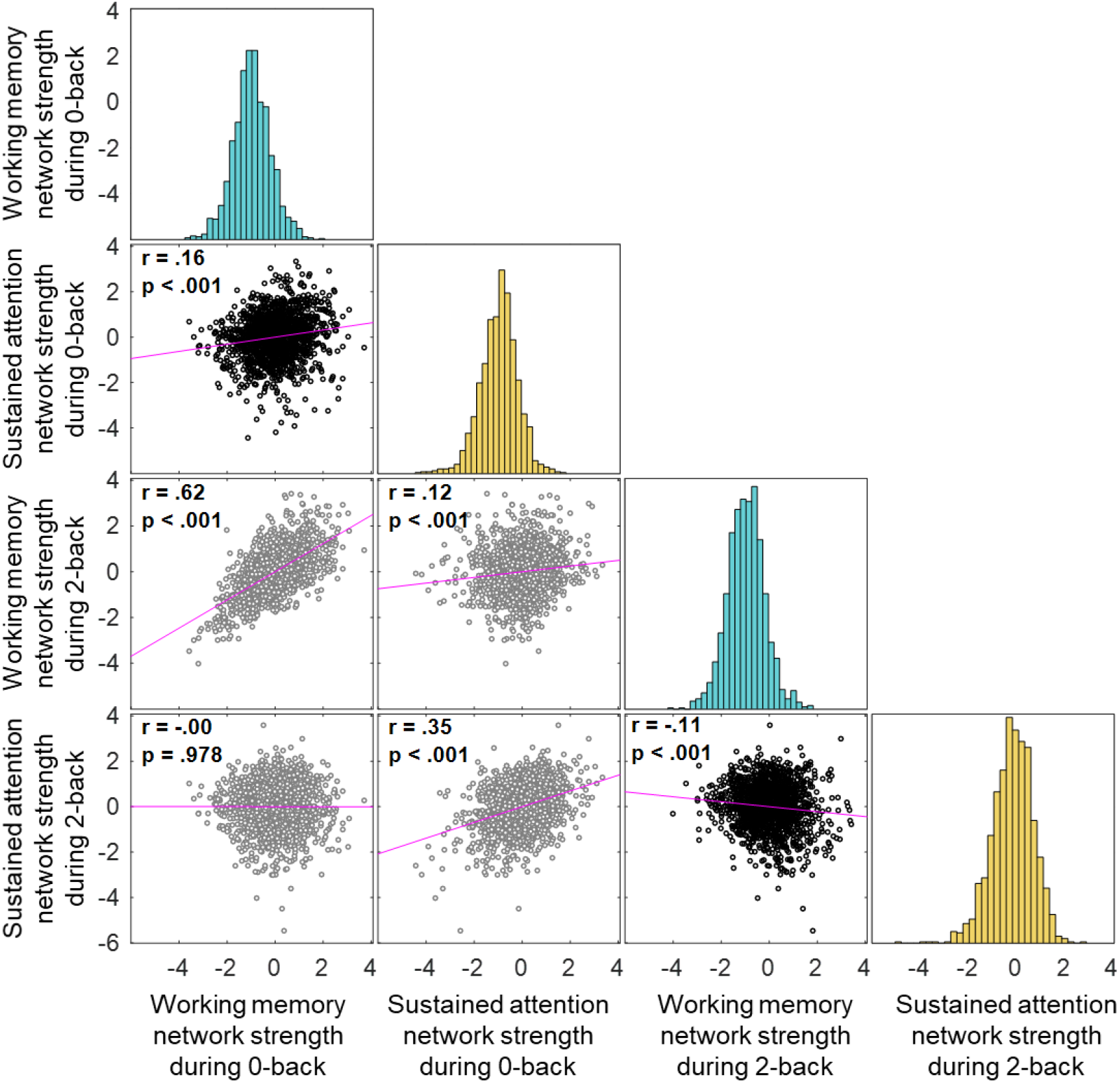
Network strengths across the participants and tasks. Correlations between predictive networks strength values across the participants in the 2-back task and 0-back tasks. Task-congruent relationships are shown in black scatterplots.

### 1.1.2. Neuromarkers differentially predict sustained attention and working memory abilities

After confirming that the sustained attention and working memory networks are separable in children, we asked whether they generalize to *specifically* predict these abilities in children. To answer this question, we related adult sustained attention and working memory network strength values to task performance during 0-back and 2-back task blocks across the 9-11-year-old participants. Again, we predicted that the sustained attention model would capture 0-back performance whereas the working memory model would capture 2-back performance.

Supporting our hypothesis, we found that strength of the adult sustained attention network predicted 0-back performance (*r* = .19, *p*_adj_ < .001) and strength of the adult working memory network predicted 2-back performance (*r* = .13, *p*_adj_ < .001) in the preadolescent youth. This external validation demonstrates cross-dataset and cross-age generalizability of the sustained attention and working memory connectome-based predictive models (Figure 4) and suggests that the functional connectivity features that predict individual differences in sustained attentional and working memory abilities in adults are present and predictive in 9–11-year-olds. These correlations are robust to the application of post-stratification weights (to account for differences between the full ABCD sample and nationally representative socio-demographics) and non-participation weights (to account for differences between the full sample and the subset that completed the *n*-back task with low head motion; see *Methods*).

**Figure 4.**
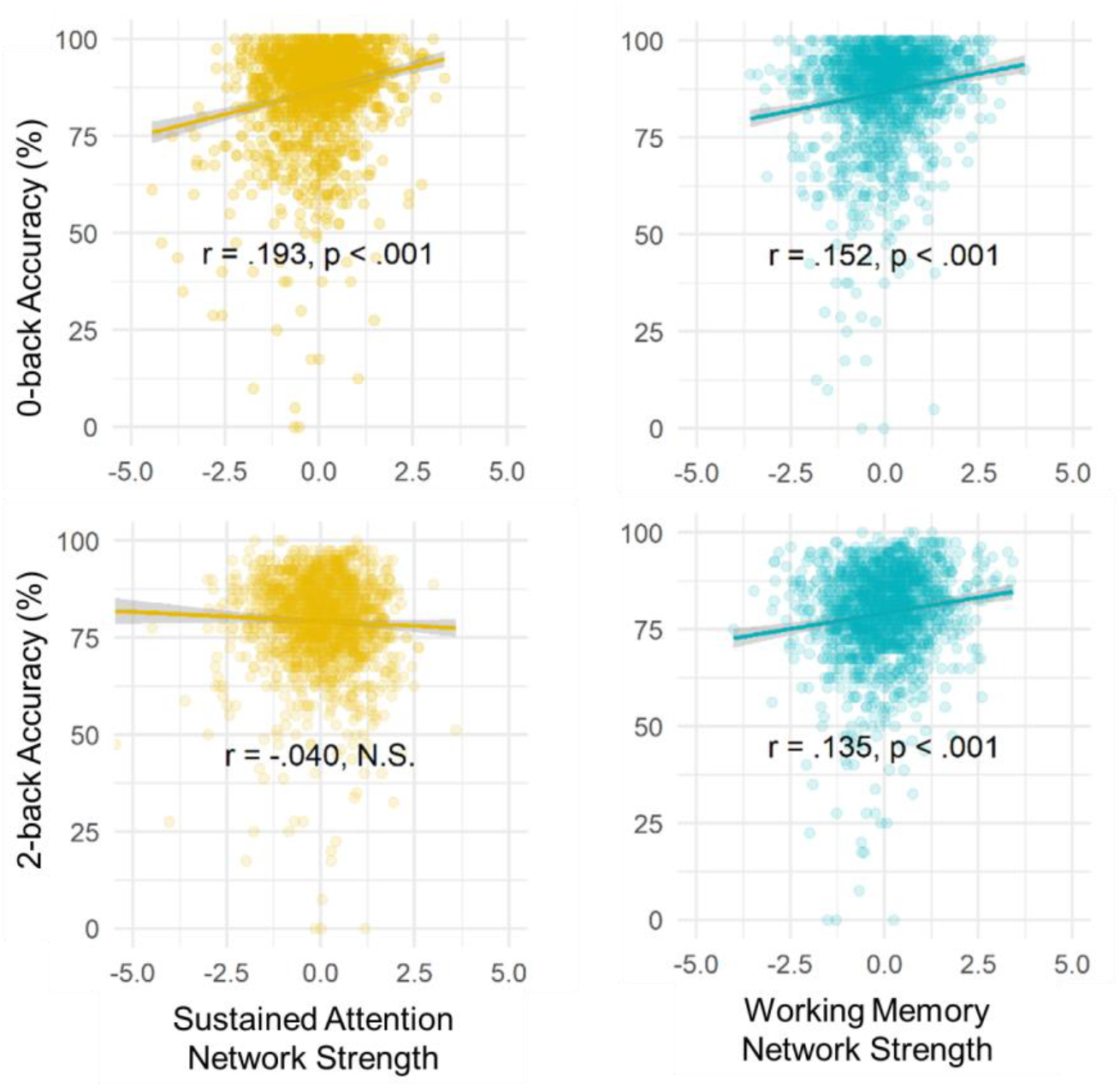
Strength of adult SA and WM networks in preadolescents predict respective task performance. Correlations between sustained attention (left, golden) and working memory (right, blue) network strength and children’s 0-back (top) and 2-back (bottom) task performance.

To assess model specificity, we compared the predictive power of the sustained attention and working memory networks for 0-back and 2-back accuracy. Sustained attention network strength was not significantly related to 2-back accuracy (*r* = –.04, *p* = .11). This correlation was significantly weaker than the correlation between working memory network strength and 2-back accuracy (William’s *t* [test of difference between two dependent correlations sharing one variable] = –4.65, *p* < .001). Thus, the working memory network was a better predictor of performance on the high-working-memory load 2-back task than the sustained attention network. We did not observe this dissociation for the 0-back task accuracy. Instead, working memory network strength predicted 0-back accuracy (*r* = .15, *p*_adj_ < .001), and this correlation was numerically but not significantly lower than the correlation between sustained attention network strength and 0-back accuracy (William’s *t* = –1.27, *p* = .20).

Strength in the sustained attention and working memory networks was correlated across children (*r* = .16, *p* < .001 during 0-back; *r* = –.11, *p* < .001 during 2-back; Figure 3), and performance in 0-back and 2-back tasks are typically correlated across individuals (*r* = .62, *p* < .001 in the current sample of 1,545 children). Thus, it is important to further assess the unique contributions of the sustained attention and working memory networks to 0-back and 2-back task performance. To this end, we included both sustained attention and working memory network strength in a regression model to predict either 0-back or 2-back accuracy (Table 1). The regression also included age, sex, and remaining head motion (after exclusion, see *Methods*) as covariates, as well as random intercepts for data collection sites. Echoing the correlation results, sustained attention network strength predicted 0-back accuracy better than chance (β = 0.16, *p* < .001) and better than it predicted 2-back performance (*p* < .001 based on bootstrapped distribution of the difference between β coefficients). In contrast, working memory network strength predicted 0-back and 2-back accuracy above chance but equally well (β = 0.11 and 0.10, respectively; *p* values < .001). Therefore, we found partial support for the specificity of the models, such that the sustained attention network predicts 0-back accuracy better than it predicts 2-back accuracy, whereas the working memory network predicts both 0-back and 2-back accuracy.

**Table 1.** Strengths of both SA and WM networks as predictors of task performance. Individual differences in sustained attention and working memory network strength are differentially related to individual differences in children’s 0-back (left) and 2-back (right) performance, respectively. “Motion” is mean frame-to-frame displacement during 0-back (left) or 2-back (right) blocks. “Sustained Attention” and “Working Memory” are mean sustained attention network strength and mean working memory network strength during 0-back (left) or 2-back (right) blocks. Marginal and conditional R^2^ statistics estimate fixed-effects R^2^ and total (i.e., fixed + random effects) R^2^, respectively, based on Nakagawa et al. (2017).

| *Predictors* | **0-back Accuracy** |  |  |  | **2-back Accuracy** |  |  |  |
| --- | --- | --- | --- | --- | --- | --- | --- | --- |
|  | *Beta* | *CI* | *t* | *p* | *Beta* | *CI* | *t* | *p* |
| *(Intercept)* | 0.04 | [-0.07, 0.12] | 0.67 | 0.513 | 0.13 | [0.02, 0.24] | 2.36 | 0.026 |
| *Age* | 0.14 | [0.09, 0.19] | 5.56 | **<0.001** | 0.15 | [0.11, 0.20] | 6.21 | **<0.001** |
| *Female* | -0.08 | [-0.18, 0.02] | -1.65 | 0.099 | -0.25 | [-0.34, -0.15] | -5.03 | **<0.001** |
| *Motion* | -0.04 | [-0.09, 0.01] | -1.59 | 0.112 | -0.13 | [-0.18, -0.08] | -5.28 | **<0.001** |
| Sustained Attention Network Strength | 0.16 | [0.11, 0.21] | 6.35 | **<0.001** | -0.04 | [-0.09, 0.00] | -1.77 | 0.078 |
| Working Memory Network Strength | 0.11 | [0.06, 0.16] | 4.19 | **<0.001** | 0.10 | [0.05, 0.15] | 3.89 | **<0.001** |
| **Random Effects (Site)** |  |  |  |  |  |  |  |  |
| ICC |  | 0.03<sub>site</sub> |  |  |  | 0.03<sub>site</sub> |  |  |
| Observations |  | 1545 |  |  |  | 1545 |  |  |
| Marginal R<sup>2</sup> / Conditional R<sup>2</sup> |  | 0.074 / 0.102 |  |  |  | 0.079 / 0.105 |  |  |

Two factors may contribute to the lack of specificity of the working memory connecotme-based model. First, the 0-back task does require memory for the target image introduced at the start of each 0-back block, and thus is a low-load rather than a no-load task. Second, the adult working memory model was originally defined to predict individual differences in 2-back task performance from adult connectomes comprised of both 0-back and 2-back fMRI data in the HCP sample (Avery et al., 2020), potentially increasing its sensitivity to *n*-back task performance overall. Future work assessing the sustained attention and working memory models’ generalizability to different datasets and behavioral measures will further inform their sensitivity and specificity.

**Study 1.2. Tracking changes in sustained attention and working memory over time**. Do the adult connectome-based models of sustained attention and working memory also vary with children’s performance fluctuations? To test this, we examined relationships between block-to-block fluctuations in network strength and block-to-block fluctuations in task performance within-participants. We also investigated whether changes in network strength and behavior were driven by stimulus types or were more spontaneous.

Mixed-effects block-level regressions with random intercepts for participants (Table 2) showed that block-by-block changes in sustained attention network strength tracked block-to-block fluctuations in 0-back accuracy (β = .07, p < .001) and block-to-block fluctuations in working memory network strength tracked block-by-block 2-back accuracy (β = .04, *p* < .001). These results were consistent with our predictions in Study 1.1. Again, demonstrating partial specificity, adult sustained attention network strength did not significantly track 2-back accuracy (β = –.01, *p* = .015) whereas adult working memory network strength did track 0-back accuracy (β = .05, *p* < .001) in youth.

**Table 2.** Block-by block networks strength and task performance. Block-by-block changes in sustained attention and working memory networks strength values are differentially related to block level 0-back and 2-back performance in children, respectively.

| 0-back Accuracy |  |  |  |  | 2-back Accuracy |  |  |  |
| --- | --- | --- | --- | --- | --- | --- | --- | --- |
| Predictors | Beta | CI | t | p | Beta | CI | t | p |
| (Intercept) | 0.07 | [0.02, 0.12] | 2.93 | **0.003** | 0.03 | [-0.01, 0.08] | 1.37 | 0.172 |
| Block Motion | -0.03 | [-0.05, -0.01] | -3.06 | **0.002** | -0.02 | [-0.04, -0.00] | -2.26 | **0.024** |
| Block Type: |  |  |  |  |  |  |  |  |
| Negative Face | -0.13 | [-0.18, -0.09] | -5.74 | **<0.001** | -0.00 | [-0.05, 0.04] | -0.16 | 0.876 |
| Positive Face | 0.01 | [-0.03, 0.06] | 0.64 | 0.522 | -0.03 | [-0.07, 0.02] | -1.16 | 0.245 |
| Place | -0.22 | [-0.26, -0.17] | -9.51 | **<0.001** | -0.36 | [-0.40, -0.31] | -15.60 | **<0.001** |
| Run 2 – Run 1 | 0.02 | [-0.02, 0.05] | 0.80 | 0.424 | 0.15 | [0.11, 0.18] | 7.75 | **<0.001** |
| Block Sustained<br>Attention Network<br>Strength | 0.07 | [0.05, 0.09] | 7.37 | **<0.001** | -0.01 | [-0.03, 0.00] | -1.46 | 0.145 |
| Block Working<br>Memory Network<br>Strength | 0.05 | [0.03, 0.07] | 5.56 | **<0.001** | 0.04 | [0.02, 0.06] | 3.83 | **<0.001** |
| **Random Effects**<br>**(Subject)** |  |  |  |  |  |  |  |  |
| ICC | 0.40<sub>subs</sub> |  |  |  | 0.39<sub>subs</sub> |  |  |  |
| Observations<br>(blocks) | 9176 |  |  |  | 9176 |  |  |  |
| Marginal R<sup>2</sup> /<br>Conditional R<sup>2</sup> | 0.019 / 0.412 |  |  |  | 0.030 / 0.411 |  |  |  |

The observed relationships between functional network strength and task accuracy are above and beyond the variance in block-by-block *n*-back accuracy explained by potential practice effects (i.e., run 2 vs. run 1) or stimulus type (i.e., positive vs. neutral faces, negative vs. neutral faces, and places vs. neutral faces; see Supplementary results: *Stimulus type does not explain within-subject relationships between network strength and behavior*) because these potential sources of variance are included as covariates in the regression model. Despite the numerically small effect sizes, it is noteworthy that the strength of sustained attention and working memory networks— developed in completely independent datasets to predict *individual differences* in <u>adults</u>—track *within-person fluctuations* in 0-back and 2-back accuracy in <u>children</u>.

**Study 1.3. Working memory network strength predicts subsequent memory**. Do youth with functional connectivity signatures of stronger sustained attention and/or working memory function during memory encoding show better later visual recognition memory? To investigate this question, we measured the relationship between recognition memory performance for *n*-back task stimuli to individual differences in networks strength values during the *n*-back task.

Recognition memory was assessed after scanning sessions. The *n*-back recognition memory test included 48 “old” stimuli (which has been presented during the *n*-back task) and 48 “new” stimuli (which has not been presented), with 12 images each of happy, fearful, and neutral faces as well as places. Participants were asked to rate each picture as either “old” or “new.” Memory performance was measured as the discrimination index (*d’*) based on all stimuli. Recognition memory d’ was related to strength in the sustained attention and working memory networks averaged across all blocks (i.e., both 0-back and 2-back blocks). Results revealed that strength in the working memory (*r* = .12, *p*_adj_ < .001), but not the sustained attention (*r* = .01, *p* = .55), network predicted subsequent recognition memory (supplementary Figure S3).

Unsurprisingly, in-scanner *n*-back performance was correlated with subsequent recognition memory performance across participants (*r* = .31, *p* < .001)^1^. Nevertheless, the relationship between working memory network strength and subsequent recognition memory remained significant even when in-scanner *n*-back performance accuracy was included in the regression model as a predictor (β = 0.04, *p* = .025; Table 3) along with the age, sex, and residual head motion. Thus, the variance in recognition memory performance captured by the working memory network is not fully accounted for by in-scanner task performance. This result highlights the unique contribution of the connectivity-based measures to long-term memory predictions.

**Table 3.** Working memory network strength during in-scanner *n*-back task performance is related to subsequent recognition memory for *n*-back task stimuli after adjusting for nuisance variables and even *n*-back performance itself.

| *Predictors* | Recognition Memory $d'$ | | | |
| --- | --- | --- | --- | --- |
|  | *Beta* | *CI* | *t* | *p* |
| (Intercept) | 0.73 | [0.67, 0.79] | 23.31 | **<0.001** |
| Age | 0.06 | [0.03, 0.09] | 3.87 | **<0.001** |
| Female | -0.02 | [-0.08, 0.04] | -0.65 | 0.514 |
| Motion | 0.01 | [-0.02, 0.04] | 0.74 | 0.459 |
| *n*-back Accuracy | 0.18 | [0.14, 0.21] | 10.62 | **<0.001** |
| Working Memory Network Strength | 0.04 | [0.00, 0.07] | 2.24 | **0.025** |
| Sustained Attention Network Strength | -0.02 | [-0.05, 0.01] | -1.34 | 0.179 |
| **Random Effects** |  |  |  |  |
| **(Site)** |  |  |  |  |
| ICC | 0.02<sub>site</sub> |  |  |  |
| Observations<br>(subjects) | 1489 |  |  |  |
| Marginal $R^2$ / Conditional $R^2$ | 0.104 / 0.120 | | | |

**Study 2 overview.** In Study 2 we directly compared the adult and preadolescent brain networks that support sustained attention and working memory. In **Study 2.1**, we benchmarked the performance of the predefined adult network models in two ways to assess the effects of cross-dataset, cross-task, and cross-age generalization on models’ predictive power. In **Study 2.2** we asked how networks predicting sustained attention and working memory are differently configured in children and adults.

**Study 2.1 Benchmarking the predictive power of the adult sustained attention and working memory network models.** Although the adult sustained attention and working memory network models successfully generalized to predict inter- and intra-individual differences in *n*-back task performance in the ABCD sample, effect sizes were modest. In Study 2.1, we benchmarked these effect sizes in two ways. First, we asked how close the predictive power of the adult sustained attention and working memory models came to a model of general cognitive ability trained in the ABCD sample itself. (We did not train separate ABCD-specific sustained attention and working memory network models because the ABCD task battery does not include an out-of-scanner sustained attention measure.) Second, we asked how close the predictive power of the sustained attention model in Study 1 came to a theoretical maximum for the 0-back and 2-back tasks by applying the same model to data from the high-quality adult HCP sample. (We could not fairly perform this analysis with the working memory model because it was defined using HCP data.)

### 2.1.1. Building a development-specific connectome-based predictive model

To ask how close the predictive power of the adult sustained attention and working memory models came to that of a ‘premature’ network predictor of general cognitive ability trained in the ABCD sample itself, we defined a new connectome-based model—the **cognitive composite** network model—using leave-one-site-out cross-validation in the ABCD Study dataset (see *Methods*). The cognitive composite network model was defined to predict children’s average performance on five out-of-scanner NIH Toolbox tasks (i.e., their “cognitive composite” score; see *Methods*) because NIH Toolbox data were collected outside the scanner and the cognitive composite score was similarly correlated with 0-back accuracy (*r* = .31, *p* < .001) and 2-back accuracy (*r* = .33, *p* < .001). Thus, it was fair to use the cognitive composite network model to benchmark the predictive power of both the sustained attention networks and working memory network models. (In other words, a model built to predict cognitive composite scores would not be biased at the outset to better predict 0-back or 2-back accuracy.)

Demonstrating its utility for this analysis, the cognitive composite model successfully predicted cognitive composite scores in left-out ABCD Study sites (*r* = .295, *p* < .001 across all sites; see Supplementary Figures S4 and S5). The youth cognitive composite network (averaged over all the site-wise models and binarized at a threshold of 0.5) included edges spanning widespread cortical and subcortical-cerebellar areas (Figure 5).

**Figure 5.**
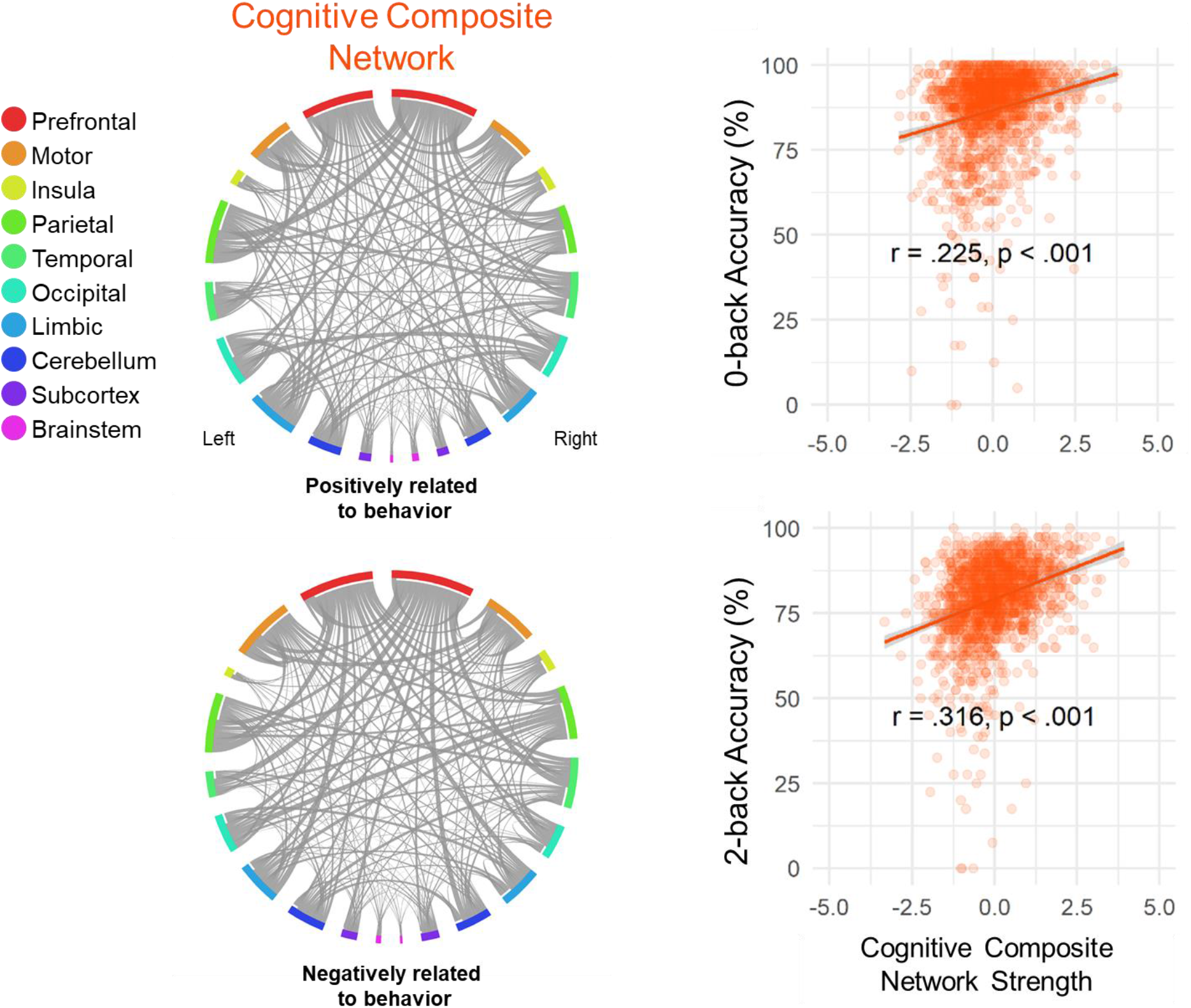
**Left:** The youth-defined cognitive composite network averaged over all the ABCD site iterations (binarized at a 0.5 threshold). **Right:** Cognitive composite network strength in 0-back and 2-back task blocks predict 0-back accuracy and 2-back accuracy across the ABCD sample, respectively.

Cognitive composite network strength during 0-back task performance predicted 0-back accuracy (*r* = .23; *p* < .001) and strength during 2-back task performance predicted 2-back accuracy (*r* = .32, *p* < .001) in children from left-out sites (Figure 5; Supplementary Table S1). This youth cognitive composite model significantly outperformed the adult working memory model for predicting 2-back accuracy (β = .27 vs. β = .10, *p* < .001 from bootstrapping). Surprisingly, however, the adult sustained attention model’s prediction of 0-back accuracy in youth was comparable with that of this ABCD-specific cognitive composite model (β = .16 versus β = .19, *p* = .242, N.S.). Furthermore, including both adult sustained attention and youth cognitive composite network strength of youths in a regression model to predict their 0-back accuracy results in comparable β coefficients for each (.18 and .21, respectively; Supplementary Table S2).

Finally, intra-individual differences analyses revealed that block-to-block changes in the strength of the youth cognitive composite network tracked block-by-block changes in both 0-back and 2-back accuracy. Echoing the block-by-block results observed with the adult network models in Study 1.2 (Table 2; sustained attention network tracking 0-back accuracy: β = .07; working memory network tracking 2-back accuracy: β = .05), the effects were significant but subtle (Supplementary Table S3, cognitive composite network tracking 0-back accuracy: β = .05; cognitive composite network tracking 2-back accuracy: β = .08).

### 2.1.2 Predicting n-back accuracy in adults

Compared to the original studies in which these networks were identified, both the sustained attention (Rosenberg et al., 2016b) and working memory (Avery et al., 2020) models show lower predictive power in the current study than they did in adults. This could arise for many reasons, including those related to developmental change (i.e., differences between adults and children) and unrelated to development (e.g., differences in scan sites and parameters and differences in the to-be-predicted behavioral task).

We used the Human Connectome Project (HCP) dataset to assess the degree to which differences *unrelated* to developmental change—scan site and parameters and task differences—impacted the predictive power of the sustained attention model. To do so, we replicated the analyses in Studies 1.1 and 1.2 with *n*-back task HCP data and compared model performance to that achieved in the ABCD dataset. A result that the model predicted adults’ 0-back accuracy better than it predicted children’s would suggest that adult models do not well-capture the functional networks underlying sustained attention performance at age 9–11 and/or that predictive power was lower in the ABCD sample because of data quality. On the other hand, a result that the model did *not* predict adults’ 0-back accuracy better than it predicted children’s would suggest that adult models *do* capture the functional networks underlying sustained attention at age 9–11. In this case, predictive power may have been lower in the ABCD sample (than in other adult datasets; e.g., Rosenberg et al., 2020) because of site- or scanner-related differences or differences in the to-be-predicted behavioral measure of sustained attention (gradCPT *d’* in Rosenberg et al., 2016b and 2020 vs. 0-back accuracy in the ABCD and HCP samples).

HCP analyses included behavioral and fMRI data from 754 adults (405 female, 22-25 years old: 174, 26-30 years old: 321, 31-35 years old: 249, and 36+ years old: 10; see *Methods*). We applied the adult sustained attention network mask to functional connectivity patterns of novel adults from HCP calculated during 0-back and 2-back blocks of the *n*-back task, and related network strength to task performance both across and within subjects. (Again we did not apply the adult working memory network mask to HCP data because it was previously defined in this sample; Avery et al., 2020.)

Demonstrating cross-dataset validity—and replicating the pattern of results observed in the ABCD sample—the adult sustained attention network predicted individual differences in novel adults in 0-back accuracy (*r* = .17, *p* < .001) but not 2-back accuracy (*r* = .07, *p* = .07; Supplementary Figure S6), with the former correlation being significantly larger than the latter (Steiger’s Z [test for the difference between two dependent correlations with different variables] = 2.29, *p* = .02). Results were consistent after adjusting for age, sex, and remaining head motion covariates (see Supplementary Table S4), and the β coefficient was significantly larger for 0-back than 2-back accuracy (β = .16 vs. .07; *p* = .034 from a bootstrap test). Mixed-effects regressions showed that, within-subject, block-by-block changes in sustained attention network strength tracked block-by-block changes in 0-back accuracy (β = .08, *p* < .001; Supplementary Table S5). Thus, the adult sustained attention network generalized to a novel sample of adults to predict 0-back, but not 2-back, task performance, which similar to what we had also observed in the ABCD sample (Study 1).

The predictive power of the adult sustained attention network was numerically similar for children’s and novel adults’ 0-back task performance (between-subjects: ABCD *r* = .19 golden line in Figure 6, HCP *r* = .17 purple line in Figure 6). This was also true for tracking changes in performance within subjects (ABCD β = .07, HCP β = .08). This suggests that the sustained attention network model captures children’s 0-back (i.e., sustained attention) performance *just as well* as it captures adults’.

**Figure 6.**
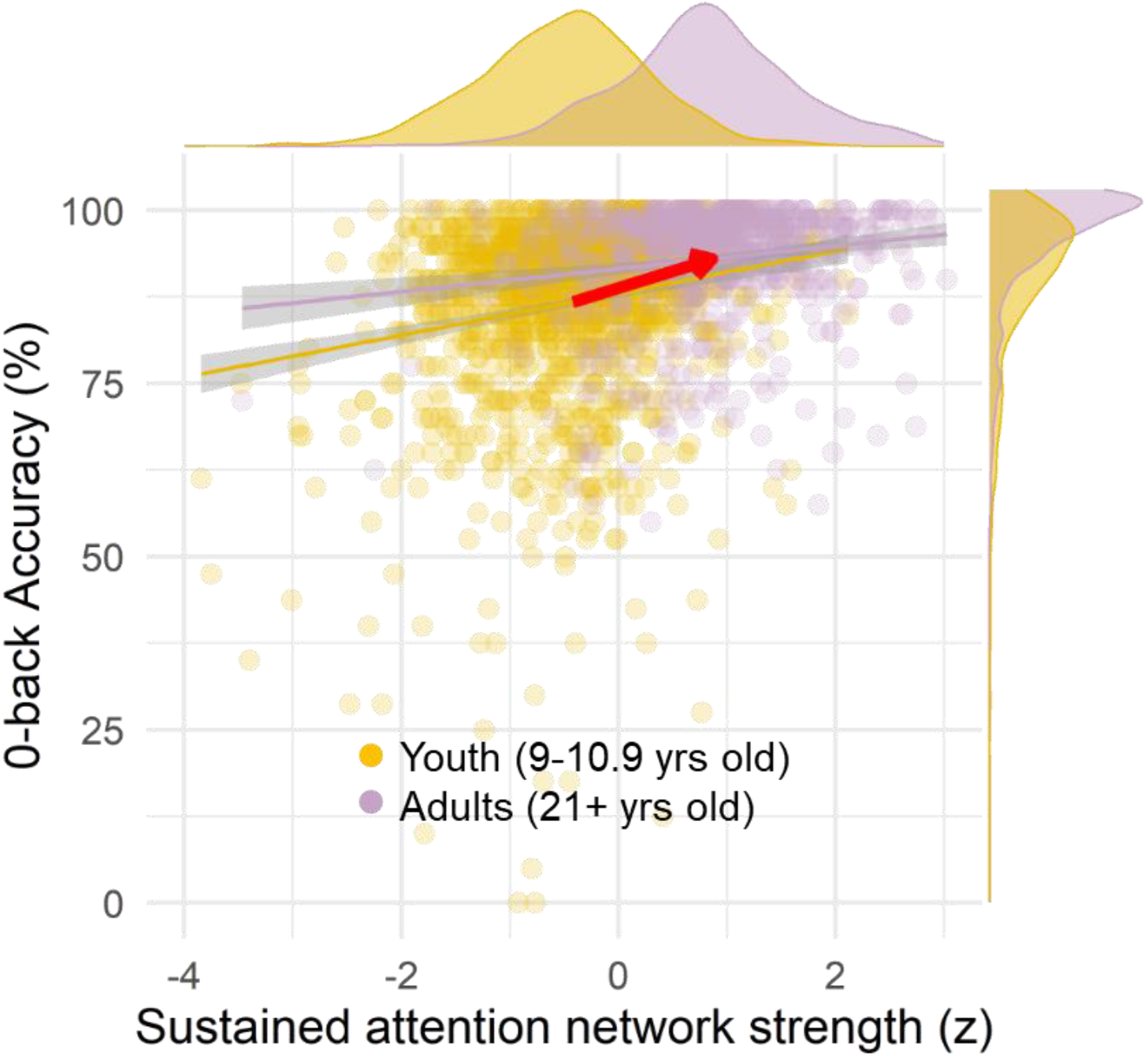
The strength of the adult SA network predicts 0-back accuracy in youth and novel adults. Even though the discriminability of individuals’ task performance is not significantly different within each dataset, i.e. there is no significant difference between the correlations (ABCD = gold, r = .19; HCP = violet, r = .17; difference between correlations Z = .46, p =.456), the mean performance and mean network strength are both significantly larger in adults.

Notably, the behavioral performance (mean 0-back accuracy) for adults is higher than youth by 6.2%±0.8% (in Figure 6 follow the red arrow along the y axis; Welch two sample t(2130.6) = 14.3, p < .001), suggesting sustained attention ability is on average stronger in adults than youth. There are multiple possibilities that could explain this. One is that the preadolescent sustained attention network is a pre-mature version of the adult sustained attention network and predicts individual differences in youths similar to adults but is on average expressed less strongly at 9–11 years old. For example, in Figure 6 the SA network expressed in the functional connectome of the adults is stronger than the 9–11year-olds (follow the red arrow along the x axis; Welch two sample t(1448.9) = 35.8, p < .001). Functional connectivity matrices were z-scored <u>within</u> each participant when used to quantify the sustained attention network strength values for this analysis, but this does not guarantee that scanner differences between HCP and ABCD studies do not bias the group-average network strength values (e.g. spatially non-homogenous differences between HCP and ABCD scans unrelated to development). Therefore, the current analyses cannot verify this explanation until longitudinal data from the same youths are processed, allowing mediation tests.

It is also possible that different subcomponents of the adult sustained attention network predict task performance in adults and youth. To investigate this possibility, we “computationally lesioned” edges with at least one node in each of ten macroscale brain regions from the sustained attention model. We compared the effects of computational lesioning on the prediction of 0-back accuracy in the HCP and ABCD samples by comparing the ΔR^2^ in lesioned vs. the full sustained attention network strength models. We found that lesioning the prefrontal and temporal lobes decreased prediction power more in adults than it did in children (*p* = .015 for the prefrontal and *p* = .007 for the temporal lobe based on bootstrap distribution of ΔR^2^ values). Lesioning the subcortex, on the other hand, decreased prediction power more in children than it did in adults (*p* = .009). Therefore, the full sustained attention network generalized equally well to adults and youth, though features within this network may contribute differentially to prediction at different ages.

Together Studies 2.1.1 and 2.1.2 demonstrate that the adult sustained attention model captures youth’s individual differences and fluctuations in attention just as well as it captures novel adults’—and it is no worse at predicting attention in youth than is a youth-specific model of cognition defined in the ABCD dataset itself. Furthermore, the adult working memory model captures general aspects of attention and memory in youth, and is outperformed by a youth model of general cognition, potentially suggesting less consistency in the functional architecture of working memory vs. sustained attention from age 9–11 to young adulthood.

**Study 2.2 Differences between adult and preadolescent networks.** Next, we asked, in a data-driven manner, how the networks are differently configured in children and adults and how these differences relate to sustained attention and working memory performance. This analysis is distinct from the lesion analysis in Study 2.1.2 in that no predefined networks are being used to restrict the establishment of brain-behavior relationship differences between youth and adults.

To this end, we combined all whole-brain functional connectomes from the ABCD and HCP datasets. We then applied a Partial Least Squares (PLS) regression analysis (McIntosh et al., 2004; Krishnan et al., 2011) that finds the linear combination of functional connections that maximally explains the age X 0-back performance covariance in these data (see *Methods*). This analysis revealed the multivariate patterns of functional connectivity that robustly covaried with latent variables of age (child vs. adult) and sustained attention performance simultaneously. This process was then repeated for the 2-back performance to find the common vs. age-specific functional connections that predict WM performance in adults and children.

Results of this analysis reveal two kinds of network configurations: one predicts cognitive performance *across* both age groups while the other predicts performance *specifically* in youth or adults. In other words, PLS finds a pair of orthogonal latent variables, one for brain-behavior relationships that are common for youths and adults and one for those that differ between youths and adults. Either the similarity or difference latent variable can emerge as dominant. None, one, or both latent variables can be statistically significant with reliable loadings on connectivity and performance variables. Our expectation was that the commonalities would be greater than the differences based on the results of the previous studies showing comparable fit of the adult networks to the preadolescent brain. Critically, this analysis includes full connectomes and does not “constrain” functional networks to any predefined predictive models.

The sustained attention PLS analysis primarily showed functional connectivity patterns that predicted better attentional performance in both ABCD youth and HCP adults (Figure 7 **top left**). In contrast, the second latent variable revealed the connections whose relationship to sustained attention performance was dependent on the age group, showing connections that are robustly related to better SA performance in youths and worse performance for the adults (Figure 7 **bottom left**). Specifically, the primary LV consisting of 3452 significant edges shows a pattern of connections that has a large positive correlation with 0-back accuracy in both youths (r = .51, CI = [.43, .59]) and adults (r = .63, CI = [.59, .67]). In contrast, the second LV consisting of 165 significant edges shows a pattern of connections that separate out the youths and adults in their task performance, as it is positively correlated (r = .79, CI = [.74, .83]) with 0-back accuracy in preadolescents while negatively in adults (r= -.40, CI = [-.50, -.28]).

**Figure 7.**
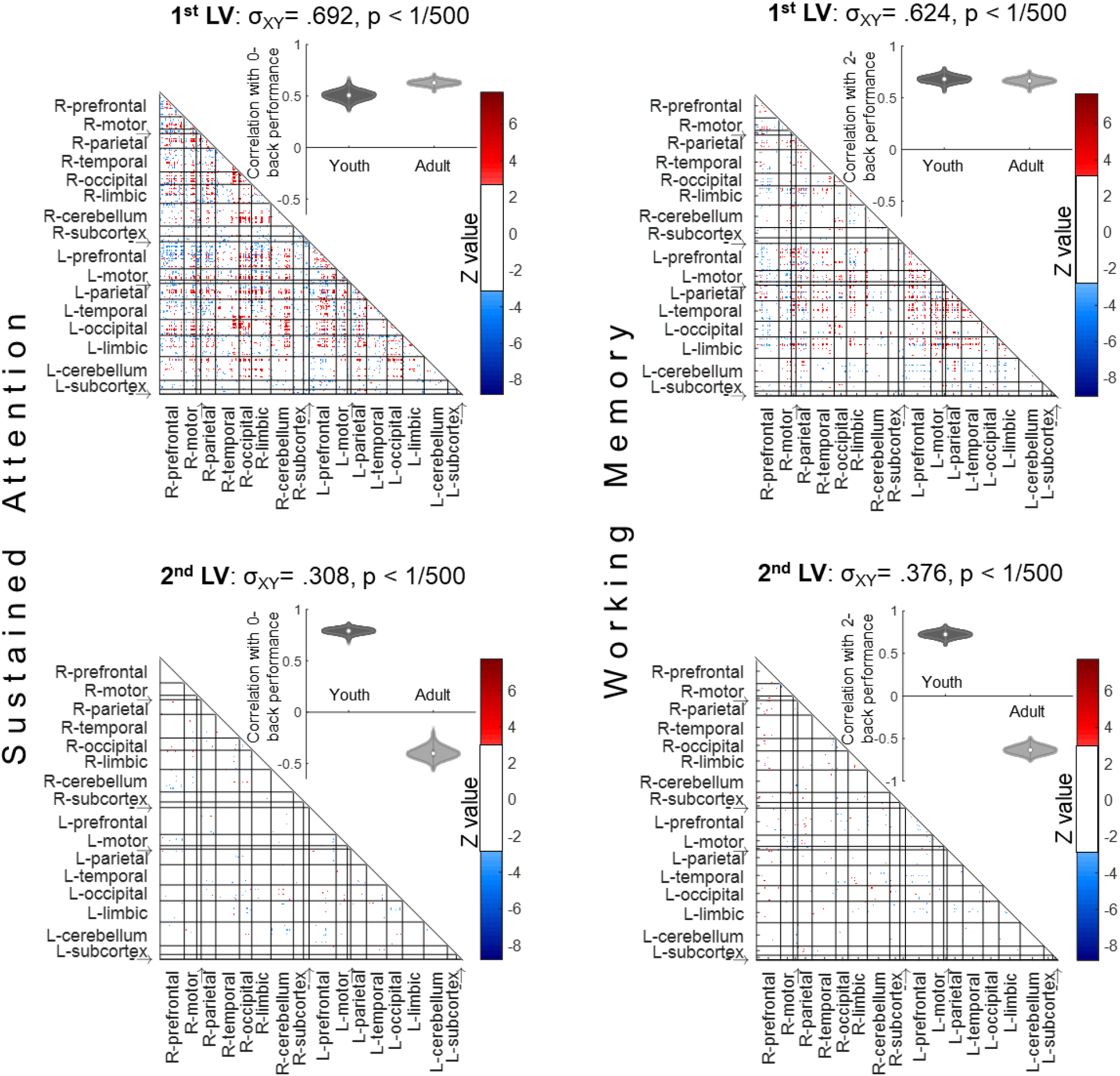
The data-driven latent variables (LV) for the sustained attention (left column) and working memory (right column) PLS regressions. **Left column:** PLS relating functional connectivity to sustained attention performance and age group successfully identified functional connectivity patterns related to better attentional performance in both ABCD youth and HCP adults (1^st^ LV) as well as connections differentially related to performance in youth and adults (2^nd^ LV). **Right column:** PLS relating functional connectivity to working memory performance and age group successfully identifies functional connectivity patterns related to better working memory performance in both youth and adults (1^st^ LV), as well as connections differentially related to performance in youth and adults (2^nd^ LV). P-values are calculated from 500 permutations and LV weights are calculated from 500 bootstraps in each PLS; significant connections are those that have bootstrap ratios (Z) above +3 or below −3. We randomly selected 754 of the ABCD sample in each PLS to make the age group sizes balanced (HCP has n = 754) and the presented figure is averaged over 200 bootstrapped balanced samples. Using the full 1545 ABCD participants instead of randomly balanced group sizes results in very similar PLS latent variables. Anatomical labels: R and L refer to right and left hemispheres; ‘→’ is insula and ‘-→’ is brainstem.

A similar pattern of results was observed for the working memory PLS analysis, where the first latent variable included functional connections that were related to working memory performance similarly for youth and adults (Figure 7 **top right**) whereas the second latent variable showed functional connections that related to working memory performance differentially for youth vs. adults (Figure 7 **bottom right**). The primary working memory LV consisted of 1610 significant edges and predicted higher 2-back accuracy in youth (r = .68, CI = [.62, .74]) as well as adults (r = .66, CI = [.60, .71]). The second LV consists of 224 significant edges and is related to better 2-back performance in preadolescents (r = .73, CI = [.66, .79]) but poorer performance in adults (r = -.63, CI = [-.72, -.55]).

Importantly, these PLS results are consistent with our neuromarker generalizability approach by showing the first latent variable, which represents connections that are positively related to performance for both youths and adults, is *stronger* for sustained attention than working memory (cross-block covariance^2^: attention σ_XY_ = .692; working memory σ_XY_ = .624 and these covariance scores are significantly different from one another *p* < 1/200 based on 200 bootstraps). Equivalently, latent variable 2 that represents functional connections that relate to performance differentially for youth vs. adults is *stronger* for working memory than sustained attention (attention σ_XY_ = .308; working memory σ_XY_ = .376; *p* < 1/200 for the difference between these cross-block covariances). These analyses further support the idea of more developmental change from preadolescence to adulthood in networks supporting working memory and more developmental stability from preadolescence to adulthood in networks supporting sustained attention.

## Discussion

Across two studies, we used different approaches using connectome-based predictions to reveal developmental change in the functional architecture of sustained attention and working memory between preadolescence and adulthood. The first approach utilized adult network models developed previously, and allowed us to evaluate the degree to which these same networks predicted sustained attention and working memory in preadolescence. The second approach directly compared and contrasted the functional connectivity underlying sustained attention and working memory in youth compared to adults.

In Study 1 we found that connectome-based models of sustained attention and working memory previously defined in independent samples of adults generalized to capture inter-and intra-individual differences in sustained attention and working memory in 9– 11-year-olds. In Study 2 we showed that the adult sustained attention network predicted children’s 0-back task performance just as well it predicted novel adults’, and just as well as a youth-defined network predictor of general cognitive abilities. The adult working memory model, on the other hand, predicted children’s 0-back and 2-back (i.e., low- and high-working memory load) task performance, although not as well as the development-specific model of cognitive abilities. These results suggest that distinct functional brain networks predict sustained attention and working memory in the developing brain. Furthermore, working memory network strength during the *n*-back predicted subsequent memory for the items in the recognition memory task performed later outside the scanner, even when adjusted for n-back performance. This result demonstrates that, in addition to predicting ongoing working task performance, working memory network expression predicts future long-term memory.

The current work signifies three benefits of individualized predictive modeling with fMRI—and, in particular, of validating predictive markers in multiple independent datasets. First, training and testing brain-based predictive models allows us to investigate specific versus general brain markers of cognitive processes by conducting single- or double-dissociation analyses predicting individual differences in different aspects of cognition. This can inform the extent to which different processes relate to common or distinct functional network. For example, sustained attention and working memory are highly related processes as they covary together in individual ability (Unsworth & Robison, 2020; Adam et al., 2015), and attention lapses lead to worse working memory performance (deBettencourt et al. 2018). Additionally, the ability to control attention has been proposed to play a major role in complex working memory tasks (Barrett, Tugade, & Engle, 2004; Kane, Bleckley, Conway, & Engle, 2001). However, our results suggest that networks involved in sustaining attention are not sufficient to predict differences in working memory (i.e., 2-back) performance across participants, despite predicting attentional (i.e., 0-back) performance in the same participants.

Second, we can ask whether the same networks that predict individual differences in behavior capture intra-individual change. Recent work has begun to suggest that fluctuations in large-scale functional brain networks index variance in sustained attention function (Rosenberg et al., 2020) and stimulus-unrelated thought (Kucyi et al., 2021) within individuals. In the current study, we demonstrate that block-by-block changes in sustained attention network strength generalized to tracked block-to-block fluctuations in 0-back accuracy of children, and block-to-block fluctuations in working memory network strength tracked block-by-block 2-back accuracy, above and beyond stimulus types and practice effects. This is remarkable given the relatively few volumes of data per block (30-31 TRs) and blocks per run. We also explored the *source* of changes in predictive network strength by investigating whether network fluctuations were more driven by *(a)* an individual’s cognitive/attentional state fluctuations or *(b)* properties of the task stimuli that they saw (i.e., the category of *n*-back task images). We found that associations between network and *n*-back performance fluctuations are largely independent of the effects of stimulus type on performance. With further longitudinal data, it will be possible to directly model intra-individual changes in functional connectivity patterns that covary with performance, thus assessing the similarities and differences in trait-like versus state indicators of sustained attention and working memory processes in functional connectivity space.

Third, applying predictive models to predict differences in behavior both between and within individuals allows us to assess how brain systems underlying different cognitive processes change (or remain consistent) across different time scales, from one moment, hour, and even year to the next. Although the adult neuromarkers of sustained attention and working memory both generalized to predict 9-11-year-olds’ behavior, they differed in their comparative fit to youth versus novel adults. This may reflect differential developmental effects for each of the two cognitive constructs.

For working memory, the adult network’s predictive power in the ABCD sample was smaller than it was in the external validation sample of older adults in the Avery et al., 2020 study (ABCD *r* = .14; older adults *r* = .36 [Avery et al., 2020]). Second, it was also smaller than the predictive power of the ABCD-defined cognitive composite network model (correlations with individual differences in 2-back accuracy: working memory network *r* = .14; cognitive composite network *r* = .32). These two comparisons converge to suggest the possibility of developmental changes in the functional architecture of working memory during adolescence into adulthood. Interestingly, there is a relatively large and significant overlap between the youth-defined cognitive composite network and the adult working memory network both for edges positively related to behavior (11.6% overlap of combined edges, hypergeometric *p* < .001), and edges negatively related to behavior (10.7% overlap of combined edges, *p* < .001). This overlap may reflect a common subnetwork underlying general cognitive ability in youth and working memory in adulthood.

For sustained attention, however, the cross-development results paint a different picture. The performance of the sustained attention network in predicting 0-back performance was similar for youth and adults (*r* = .19 and .17, respectively). Although this may reflect similarity in the functional architecture of sustained attention in these two age groups, it could arise from better “ground truth” prediction in adulthood disadvantaged by the relatively low variance in their 0-back accuracy (HCP SD = .081 vs. ABCD SD = .126). Making this explanation unlikely, however, the child cognitive composite network did not significantly outperform the sustained attention network in predicting 0-back accuracy (*r* = .23 vs. .19, respectively) despite the identical variance in behavioral performance (i.e., both predictions are in ABCD sample). This makes the difference in variance between adults and children a less tenable explanation for similarities in prediction and instead indicates that the sustained attention network is as informative about the functional architecture of sustained attention in youth as it is in adulthood. The result suggests consistency in this architecture from preadolescence to adulthood. Additionally, the relationship between children’s sustained attention network strength and 0-back accuracy was unchanged when adjusting for the strength of the cognitive composite network, again pointing to its unique and specific relation to sustained attention even in youth.

Although the sustained attention network model generalized equally well to youth and adult 0-back performance, the particular contributions of different anatomical regions involved in the network differed between the two populations. That is, lesioning prefrontal, temporal and subcortical regions from the networks affected predictive power differently for youth and adults. This may be related to the fact that sustained attention function does improve well into adulthood (Fortenbaugh et al., 2015), and in these samples 0-back performance was indeed higher in the HCP than the ABCD data (mean accuracy = .93 vs. .87, *t* (2130.6) = 14.3, *p* < .001). Therefore, it may be the case that the neural markers of individual differences in sustained attention are present and predictive by late childhood, but the way they are utilized to maintain focus on tasks may change through adolescence. Taking together these and the working memory results discussed earlier, we found differential developmental effects for each of the two cognitive constructs, pointing to less consistency in the functional architecture of working memory than that of sustained attention from age 9–11 to young adulthood.

Study 2.2 echoed these results by demonstrating that the relationship between functional connectivity patterns and sustained attention is more similar between youths and adults than the relationship between functional connectivity patterns and working memory. Further, this analysis revealed that most functional connections related to behavioral performance (SA or WM) are *shared* between youths and adults, as the shared component was the primary latent variable for both sustained attention and working memory in the PLS regressions. However, there are also connections, which comprise the secondary LVs, that differentiate youths from adults, i.e., connections that predict better performance in youth but worse performance in adults. Future work can use longitudinal data to ask whether these patterns reflect neurodevelopment underlying cognitive and attentional performance improvement from preadolescence through adulthood.

There are some limitations to this work. First, we rely on 0-back and 2-back performance to index sustained attention and working memory rather than more traditional tasks like a CPT and visual change detection task, as these more traditional paradigms are not included in the ABCD Study. Future work characterizing the generalizability of connectivity-based models to other tasks of attention and memory can further inform their predictive boundaries. Second, the within-participant fluctuation effects in Study 1.2 are statistically significant yet modest, and the across-participant effects in Studies 1.1 and 1.3 are small to medium. Although the current approach demonstrates the statistical significance and theoretical implications of conservative external model validation analyses, further work is needed to determine the practical significance and potential translational utility of these and other brain-based predictive models. Third, as mentioned before, in Study 2.2 since the ABCD and HCP samples are not the same cohort, it is difficult to tease apart the age-related versus scanning-related differences between the two samples in the multivariate PLS regression.

In conclusion, we found that distinct functional brain networks predict sustained attention and working memory abilities across youth, as well as changes in attentional and memory performances over time. Therefore, sustained attention and working memory are overlapping but distinguishable cognitive constructs in the pre-adolescent brain, with functional connectivity patterns of working memory changing more over adolescence and into adulthood than those of sustained attention.

## Methods

### Data

We analyzed a subset of baseline-year behavioral and fMRI data from the Adolescent Brain Cognitive Development (ABCD; Release 2.0.1). The total sample in the dataset is 11,875 children 9-11 years old from 21 sites across the US. We first excluded the participants who were scanned using Philips scanners (see *fMRI data processing*) or those without functional MRI data, resulting in 9,446 participants from 19 sites. After a visual quality check of all structural and functional scans, 4,939 of these participants had structural and at least one run of *n*-back task fMRI data that passed our visual quality check and had corresponding EPrime files containing trial-by-trial *n*-back task data. Next, we applied a frame displacement (FD) threshold of FD mean < 0.2 mm and FD max < 2 mm to remove *n*-back fMRI runs with excessive head motion, resulting in 1,839 participants. Finally, we removed *n*-back runs for which the start time of the behavioral recording file was unclear with respect to the fMRI data, or if the data were flagged for “switched box” or “*n*-back task done outside scanner”, resulting in sample size of N = 1,548. Additionally, participants from one site with only N = 3 subjects after prior exclusions were not included in the across-subjects models which include site as a random intercept factor and also the within-subject analyses for consistency. Therefore, the final sample size was N = 1545 participants 9–11 years old from 18 sites, mean age = 10.03 years old, 851 female.

### In-scanner emotional n-back task

The emotional *n*-back task in the ABCD dataset (Casey et al., 2018) includes two runs of eight blocks each with 10 trials in each block. A picture is shown in every trial and participants are told to make a response on every trial, indicating whether the picture is a “Match” or “No Match.” In each run, four blocks are 2-back task for which participants are instructed to respond “match” when the current stimulus is the same as the one shown two trials back. The other four blocks are of the 0-back task for which participants are instructed to respond “match” when the current stimulus is the same as the target image presented at the beginning of the block. At the start of each block, a 2.5 s cue indicates the task type (“2-back” or “target=” and a photo of the target stimulus; see Figure S7). A 500 ms colored fixation precedes each block instruction, to alert the child of a switch in the task condition.

Two blocks of 0-back and two blocks of 2-back contain happy faces (1 in each run), another two in each task contain fearful faces, another two contain neutral faces, and another two contain places. There are 24 unique stimuli per type presented in separate blocks, each trial is 2.5 s (2 s presentation of a stimulus, followed immediately by a 500 ms fixation cross) resulting in 160 total trials in 16 blocks of *n*-back. Four fixation blocks (15 s each) also occur in each run after of every other *n*-back block.

### Post-scan n-back recognition memory task

We analyzed data from the post-scan *n*-back recognition memory task, in which 48 “old stimuli” (previously presented during the in-scanner emotional *n*-back task) and 48 new stimuli were presented to participants. Participants were asked to rate each picture as either “Old” or “New.” Each picture was presented for 2 s followed immediately by a 1-s fixation cross. The 96 pictures shown have equal numbers of each stimulus type in the old and new stimulus sets (12 each of happy, fearful, and neutral facial expressions and places in each set).

#### Cognitive composite score

The “cognitive composite” behavioral scores were measured from each child’s average performance in five out-of-scanner NIH-Toolbox tasks: the Picture Vocabulary task, Flanker inhibitory control and attention task, Pattern Comparison processing speed task, Picture sequence memory task, and Oral Reading recognition task. These tasks were chosen because they capture a wide range of cognitive processes are the only NIH Toolbox tasks collected in subsequent ABCD data releases. This makes them (and the cognitive composite score used here) suitable for future longitudinal tracking of the general cognitive abilities of the children using the model developed from the current release. The mean of these five measures was used as the cognitive composite score rather than first principal component (PC) because the correlation between the mean and the first PC was *r* = 0.94. Therefore, we used the mean for a straightforward interpretation.

### Functional MRI data processing

Minimally preprocessed functional and structural scans for ABCD Release 2.0.1 were downloaded for all participants from the National Institutes of Mental Health data archive. Use of the data was approved by the relevant University of Chicago Institutional Review Board. Minimal preprocessing included motion correction, B0 distortion correction, gradient warping correction and resampling to an isotropic space (Hagler, 2019). Participants who were scanned on Philips brand scanners were excluded because of a known error in the phase encoding direction while converting from DICOM to NIFTI format. Next a custom modification of the FMRIPREP pipeline was run on all images. Each participant’s structural T1w scan was skull-stripped, segmented by tissue type, and then normalized to the MNI152 non-linear 6^th^ generation template: the standard MNI template included with FSL. Functional scans were then aligned and normalized to the T1w space and then to MNI space and potential confounds of interest were extracted. Next 36 confounds (Power 2012) were regressed out of the voxelwise BOLD timeseries including: global mean signal, mean cerebro-spinal fluid signal, mean white matter signal, the 6 standard affine motion parameters and their derivatives, squares, and squared derivatives. This was followed by applying a bandpass filter with a highpass cutoff of .008 Hz and a lowpass cutoff of .12 Hz via the 3dBandpass command in AFNI. Finally, the cleaned volumetric BOLD images were spatially averaged into 268 predefined parcels, including cortical, subcortical, and cerebellar regions, from the whole-brain Shen functional atlas (Shen 2013).

### Predictive network anatomy

The **sustained attention** network mask (Figure 2) was defined to predict sustained attention function using data collected from 25 adults who performed a gradual-onset continuous performance task (gradCPT; Esterman et al., 2013) during fMRI (Rosenberg et al., 2016b). Sustained attentional abilities were operationalized as participants’ sensitivity (*d’*) on the gradCPT. The same functional networks that predicted performance in the initial training sample—a “high-attention” network whose strength predicted higher *d’* scores and a “low-attention” network whose strength predicted lower *d’* scores—have generalized to independent datasets to predict performance on other attention tasks from data observed during rest and task performance. These include a stop-signal task (Rosenberg et al., 2016a), Attention Network Task (Rosenberg et al., 2018), Stroop task (Fountain-Zaragoza et al., 2019), and Sustained Attention to Response Task (Wu et al., 2019).

The sustained attention networks do not rely on canonical brain networks, such as the default mode and frontoparietal networks, to predict sustained attention. Instead, the high-and low-attention networks comprise 757 and 630 functional connections, or edges, respectively (out of 35,778 total), and span distributed cortical, subcortical, and cerebellar regions. In general, functional connections between motor cortex, occipital lobes, and cerebellum predict better sustained attention whereas functional connections between temporal and parietal regions, within the temporal lobe, and within the cerebellum predict worse attention. Computationally lesioning the high- and low-attention networks by removing connections from specific brain networks does not significantly reduce predictive model performance (Rosenberg, 2016), suggesting that the sustained attention connectome-based predictive model does not rely on individual canonical networks to achieve significant prediction.

The **working memory** network mask (Figure 2; Avery et al., 2020) was defined to predict 2-back task accuracy from data observed during 10-min *n*-back task fMRI runs (both 0-back and 2-back blocks) in the Human Connectome Project dataset (N = 502 from the S900 release). Like the sustained attention networks, the working memory networks were defined using connectome-based predictive modeling (Finn et al., 2015; Shen et al., 2017). Briefly, in this approach, a “high-working-memory” network whose strength predicted higher 2-back accuracy and a “low-working-memory” network whose strength predicted lower 2-back accuracy were identified by correlating all edges (defined with the 268-node whole-brain Shen atlas; Shen et al., 2013) with 2-back accuracy across the HCP sample and retaining the edges significantly related to performance (*p* < .01). The resulting network model predicted unseen 2-back accuracy scores in HCP sample (*r* = .36, *p* < .001) in an internal cross-validation analysis, and generalized to predict individual differences in a composite of visual and verbal memory task performance (*r* = .37, *p* < .001) from resting-state fMRI in an independent sample of 157 older adults, 109 of whom were memory-impaired (Avery et al., 2020).

The working memory networks comprise a distributed set of edges (1,674 edges in the high-working-memory network and 1,203 edges in the low-working-memory network) including frontoparietal, subcortical-cerebellar, motor, and insular edges. Additionally, default mode network (DMN) connections are included in the working memory networks, with the DMN and DMN-associated regions (including limbic, prefrontal, parietal, and temporal cortices) overrepresented in the high-working-memory network relative to the low-working-memory network (Avery et al., 2020).

### Functional connectivity measures

The stimulus onset and offset times of the first and last trial in each block of the ABCD *n*-back task data were extracted from each participant’s *n*-back EPrime file (shared in the curated MRI data folders sourcedata/func/task_events). The node-wise BOLD signal time series during each *n*-back block (30 or 31 TRs; TR = 0.8 s; ∼25 seconds from the onset of the first stimulus to the offset of the last stimulus in each block) were used to create block-wise functional connectivity matrices (FC matrices) by computing all pairwise Pearson correlations between the block-wise time series of the 268 Shen parcels. (See *Defining* n*-back blocks with a temporal lag* in the supplement for a replication analysis with different block onsets and offsets). The positive edge mask (i.e., the functional connections positively related to behavior) and negative edge mask (i.e., the functional connections negatively related to behavior) for each of the sustained attention and working memory pre-defined networks (Figure 2) were then multiplied by the FC matrices. These network masks are 268*268 trinary matrices determining if a pairwise correlation (edge) belongs to a certain predictive network with −1 or +1 or not with a 0. These are shown in Figure 2 and briefly described in the previous section. Next, the Fisher’s z-transformed correlation values in the masked FC matrices were summed (with mask weight sign) to calculate the corresponding network’s strength in the block for the participant:

Block-wise network strength = 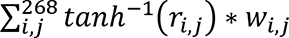

Where *r_i,j_* is the Pearson’s correlation between BOLD timesries of parcels i and j, and *w_i,j_* is the corresponding network mask value of 0, 1, or –1.

The block-wise network strengths were averaged over all 0-back blocks or all 2-back blocks for Study 1’s across-participant analyses and *z*-scored across participants. For Study 2, the measures were left at the level of blocks within each participant. In Study 4, the network strengths were averaged over all blocks (i.e., both 0-back and 2-back) for across-participant subsequent memory analysis, because the released recognition memory scores (*d’*) were from stimuli that could have been encoded during 0-back and/or 2-back task blocks, and files distinguishing the subsequent memory stimuli source were not available for this ABCD data release.

### Youth cognitive composite network

For consistency, the youth cognitive composite model was constructed using connectome-based predictive modeling (Shen et al., 2017), the same approach used to define the adult sustained attention and working memory network models. To construct the cognitive composite network mask for ABCD participants from site *k*, we retained the FC matrices (calculated from the entire *n*-back task timeseries) and cognitive composite scores of participants from all sites excluding *k*. We correlated the strength of every FC with cognitive composite score across participants in this training set. The edges positively and negatively correlated with cognitive composite score (*p* < .01) defined the masks that were applied to the block-wise functional connectivity matrices of participants from the left-out site *k* as described in the previous section. This analysis included 1536 participants because 9 participants did not have all five NIH Toolbox measurements).

### Mediation analysis

To perform the mediation analysis described in supplementary Figure S1, 2-back task performance and working memory network strength measures were mean-centered within each participant to remove individual differences and then entered into the mediation model. The model included block type (Neutral Face versus Place) as the predictor, block-wise 2-back performance as the dependent variable, block-wise working memory network strength as the mediator, and block-wise frame displacement during 2-back blocks as a covariate. The mediation was performed using the *mediation* package in R with the built-in bootstrapping option for computing *p*-values for the coefficients and the mediated effect.

### Hypergeometric cumulative distribution function

To assess whether the overlap of the edges of different predictive networks was statistically significant, we calculated the probability of the overlap being due to chance using the hypergeometric cumulative distribution function implemented in MATLAB (www.mathworks.com). The function used was *hygecdf()* computed as:

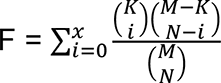

Where F is the probability of drawing up to x of a possible K items in N drawings without replacement from a group of M objects. The *p*-value for significance of overlap is then calculated as 1-F.

### Removing relatives does not change across-participant results

In our final sample (*n* = 1545) there were 82 related children (41 pairs). We repeated the across-participant analyses after randomly removing from our sample one sibling from each pair (new *n* = 1504) and found no significant differences in the results (see Supplementary Tables S6, S7, and S8). We did not repeat the block-to-block change analyses because within-participant analyses are not affected by across-participant relationships.

### Non-participation and post-stratification weights does not change across-participant results

Participant exclusion rates are high for the functional neuroimaging data in adolescents, mainly due to high attrition rate from head motion, including the current study. To correct for any biases due to not being included in the analysis relative to the demographic characteristics of the overall ABCD Study sample, we conducted sensitivity analyses to confirm if the relationships in our between-subject analyses (n = 1545) hold with non-participation (non-inclusion) weights (overall ABCD sample n = 11875). Specifically, results were re-assessed with membership in the analyses weighted by estimated non-participation weights calculated for sex, age, race/ethnicity categories, and family income and then combined with the American Census Study (ACS) to ABCD raked propensity scores in file *acspsw03.txt* of Curated Release 2.0.1 (see Heering & Berglund 2020, Dick et al 2021). The process is similar to Class et al 2019 and Stier et al 2021. In brief, inclusion/exclusion was regressed on age, sex, race/ethnicity, family income, and parental education in an elastic net regularized binary logistic regression model using glmnet in R. The logistic regression model picked the optimal tuning parameter lambda with the least cross-validation deviance in model selection. Having selected the optimal model, the non-participation weights are the inverses of the probabilities of response, conditional on being sampled. In order to compute corrected correlations between network strengths and task performances, non-participation weights capturing which individuals had available data were multiplied by the post-stratification weights. Finally, the weighted mean, weighted variance, and weighted covariance were computed as in Becker 2015, to construct the corrected correlations. No between-subject results were changed with the non-participation and post-stratification weights. Within-subject analyses are not impacted by non-participation.

### Human Connectome Project data

In Study 2, we analyzed data from the Human Connectome Project (HCP) release S1200, a multi-site consortium that collected MRI, behavioral, and demographic data from 1113 participants. Minimally preprocessed, open-access *n*-back fMRI data were downloaded from connectomeDB (https://db.humanconnectome.org/) via Amazon Web Services. The acquisition parameters and prepossessing of these data have been described in detail elsewhere (Glasser et al., 2013). Briefly, preprocessing for task data included gradient nonlinearity distortion correction, fieldmap distortion correction, realignment, and transformation to a standard space. In addition, we applied additional preprocessing steps to the minimally preprocessed task data. This included a high-pass filter of 0.001 Hz via fslmaths (Jenkinson et al., 2012), and the application of the ICA-FIX denoising procedure using the HCPpipelines (https://github.com/Washington-University/HCPpipelines) tool, which regresses out nuisance noise components effectively, similar to regressing out motion parameters and tissue type regressors (Parkes et al., 2018). The cleaned volumetric BOLD images were spatially averaged into 268 predefined parcels (Shen et al., 2013).

32 participants without sync time information files or motion regressor files for both *n*-back runs were removed from further analysis. Next, similar to the ABCD dataset, we applied a frame displacement (FD) threshold of FD mean < 0.2 mm and FD max < 2 mm to remove *n*-back fMRI runs with excessive head motion, resulting in 881 participants. Finally, we removed participants with any quality control flags from the HCP quality control process (variable QC_Issue), resulting in a final sample of 754 participants.

Functional connectivity measures for HCP data were computed as described for the ABCD data. Text files containing the timing information of the *n*-back trials were used to extract the beginning and ending of the blocks for each participant (each block ∼35 TRs; TR = 0.72 s; ∼25 seconds). A functional connectivity matrix for each block was constructed from the Pearson correlation between the BOLD signal timeseries of pairs of Shen parcels, and the sustained attention mask was applied to each block-specific FC matrix. The block-wise sustained attention network strength values were averaged over all 2-back blocks or all 0-back blocks for Study 2.1’s across-participant analyses and *z*-scored across participants. For Study 2.1’s within-participant analysis, the measures were left at the level of blocks.

### Partial Least Squares regression

In Study 2.2 we used PLS to identify the relationship between the set of connections with group-by-performance accuracy. The PLS implementation software was downloaded from Randy McIntosh’s lab at: https://www.rotman-baycrest.on.ca/index.php?section=84. In PLS, the goal of the analysis is to find weighted patterns of the original variables in the two sets (termed “latent variables” or “LVs”) that maximally co-vary with one another (McIntosh et al., 2004; Krishnan et al., 2011; examples of application for fMRI connectivity studies: Berman et al., 2014; Shen et al., 2015; Yoo et al., 2018; Kardan et al., 2019). Briefly, PLS is computed via singular value decomposition (SVD). The covariance between the two data sets X (functional connectivity matrices) and Y (age-group stacked task performances) is computed (X’Y) and is subjected to the SVD:

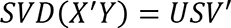

Where U and V (the right and left singular vectors) provide weights (or “saliences”) for the two sets (connectivity matrices and group-by-performance), respectively. The scalar singular value on the diagonal matrix S is proportional to the “cross-block covariance” between X and Y captured by the LV and is naturally interpreted as the effect size of this statistical association (reported as σ_XY_).

First 200 re-samples of 754 participants from the 1545 total ABCD sample were randomly selected and combined with the 754 HCP participants to make the age groups balanced. For each of these 2×754 samples, a set of 500 bootstrap samples were created by re-sampling subjects with replacement (preserving age labels) in order to determine the reliability with which each connection contributes to the overall multivariate pattern. Each new covariance matrix was subjected to SVD as before, and the singular vector weights from the resampled data were used to build a sampling distribution of the saliences from the original data set. Saliences that are highly dependent on which participants are included in the analysis will have wide distributions, therefore low reliability. For the functional connections, a single index of reliability (termed “bootstrap” ratio, or “Z_BR_”) was calculated by taking the ratio of the salience to its bootstrap estimated standard error. A Z_BR_ for a given connection is large when the connection has a large salience (i.e., makes a strong contribution to the LV) and when the bootstrap estimated standard error is small (i.e., the salience is stable across many resamplings). Here, connections with Z_BR_ > 3 or Z_BR_ < −3 were selected as showing reliable increase or decrease in functional connectivity, respectively (equivalent to p∼0.0025, 2-tailed, under normal distribution assumptions) similar to Kardan et al., 2020. In each iteration, a set of 500 covariance matrices were generated by randomly permuting condition labels for the X variables (brain set). These covariance matrices embody the null hypothesis that there is no relationship between X and Y variables. They were subjected to SVD resulting in a null distribution of singular values. The significance of the original LVs was assessed with respect to this null distribution. The *p* value was estimated as the proportion of the permuted singular values that exceed the original singular value.

## Data and code availability

All ABCD data are available at https://nda.nih.gov/edit_collection.html?id=2573. The sustained attention network masks are available at https://github.com/monicadrosenberg/Rosenberg_PNAS2020. The working memory network masks are available per request from the authors of Avery et al. (2020). HCP data are available at https://db.humanconnectome.org. Analysis scripts to generate results and figures in the manuscript will be available on https://github.com/okardan/ABCD_SA-WM upon publication.

## ABCD Acknowledgement

Data used in the preparation of this article were obtained from the Adolescent Brain Cognitive Development (ABCD) Study (abcdstudy.org), held in the NIMH Data Archive (NDA). This is a multisite, longitudinal study designed to recruit more than 10,000 children age 9-10 and follow them over 10 years into early adulthood. The ABCD Study is supported by the National Institutes of Health and additional federal partners under award numbers U01DA041022, U01DA041028, U01DA041048, U01DA041089, U01DA041106, U01DA041117, U01DA041120, U01DA041134, U01DA041148, U01DA041156, U01DA041174, U24DA041123, and U24DA041147. A full list of supporters is available at abcdstudy.org/nih-collaborators. A listing of participating sites and a complete listing of the study investigators can be found at abcdstudy.org/principal-investigators.html. ABCD consortium investigators designed and implemented the study and/or provided data but did not necessarily participate in analysis or writing of this report. This manuscript reflects the views of the authors and may not reflect the opinions or views of the NIH or ABCD consortium investigators.

The ABCD data repository grows and changes over time. The ABCD data used in this report came from NIMH Data Archive Digital Object Identifier 10.15154/1504041. DOIs can be found at nda.nih.gov/study.html?id=721.

This research also benefited from the ABCD Workshop on Brain Development and Mental Health, supported by the National Institute of Mental Health of the National Institutes of Health under Award Number R25MH120869 and by UG3-DA045251 from the National Institute of Drug Abuse.

## Acknowledgements

This research was supported by National Science Foundation BCS-2043740 (M.D.R.), National Science Foundation S&CC-1952050 (M.G.B.), National Science Foundation DGE-1746045 (K.E.S.), National Institutes of Health MH 108591 (M.M.C.), National Science Foundation BCS-1558497 (M.M.C.), the University of Chicago Micro-Metcalf Program (L.T.) and Social Sciences Division, and resources provided by the University of Chicago Research Computing Center.

## Supplementary Results

### Stimulus type does not explain within-subject relationships between network strength and behavior

As demonstrated in Study 1.2, the strength of the sustained attention and working memory networks is related to sustained attention and working memory performance within-subject even when adjusting for stimulus type. To what degree are fluctuations in behavior and functional connectivity driven by internal factors (e.g., fluctuations in intrinsic cognitive and attentional states) versus external factors (such as the characteristics of the task itself)? In other words, is the within-subject variance in *n*-back performance captured by the connectivity models driven by task design or endogenous sources? To answer this question, we examined the relationship of the *n*-back stimulus types with changes in task accuracy and changes in the sustained attention and working memory network strength. When applicable, we then conducted mediation analyses to assess the extent to which changes in networks strength mediated the effects of stimulus type on block-wise task accuracy.

We first tested whether 0-back accuracy differed as a function of stimulus type (neutral face, positive face, negative face, place). Five of the six pairwise contrasts were significant, such that 0-back accuracy was higher for neutral vs. negative faces, neutral faces vs. places, positive vs. negative faces, positive faces vs. places, and negative faces vs. places (all *p*_adj_ < .001; Tukey’s T test). Next, we calculated pairwise differences in sustained attention network strength during 0-back blocks with different stimulus types. One pairwise difference was significant, such that sustained attention network strength during place blocks was *higher* than sustained attention network strength during negative face blocks (*p*_adj_ = .026). This demonstrates that the within-subject relationship between sustained attention network strength and 0-back accuracy *cannot* be driven by stimulus type because the only stimulus-driven associations between within-subject changes in sustained attention network strength and 0-back accuracy (places vs. negative faces) are in opposite directions (Figure S1, left column).

In the 2-back task, there were three significant pairwise differences in accuracy: better performance in neutral face vs. place, positive face vs. place, and negative face vs. place blocks (all *p*_adj_ < .001). Only one pairwise difference in working memory network strength was significant, such that strength was higher during neutral face than place blocks (*p*_adj_ =.005; Figure S1, right column).

Because both 2-back accuracy and working memory network strength were higher during neutral face than place blocks, we conducted a mediation analysis to test whether the relationship between stimulus type (neutral face vs. place) and 2-back accuracy was mediated by changes in working memory network strength. Results demonstrate that changes in working memory network strength partially mediated the effect of stimulus type on block-wise 2-back accuracy (significant differences between total effect c and direct effect c’ in the mediation; c-c’ = .004, p = .004; Figure S2). The proportion of mediated effect was modest (about 1% of the total effect).

Together these findings suggest that changes in sustained attention and working memory network strength are driven to a greater degree by intrinsic fluctuations than stimulus characteristics, and that within-subject associations between network strength and task accuracy are, to a large extent, independent of stimulus type in this task.

**Figure S1.**
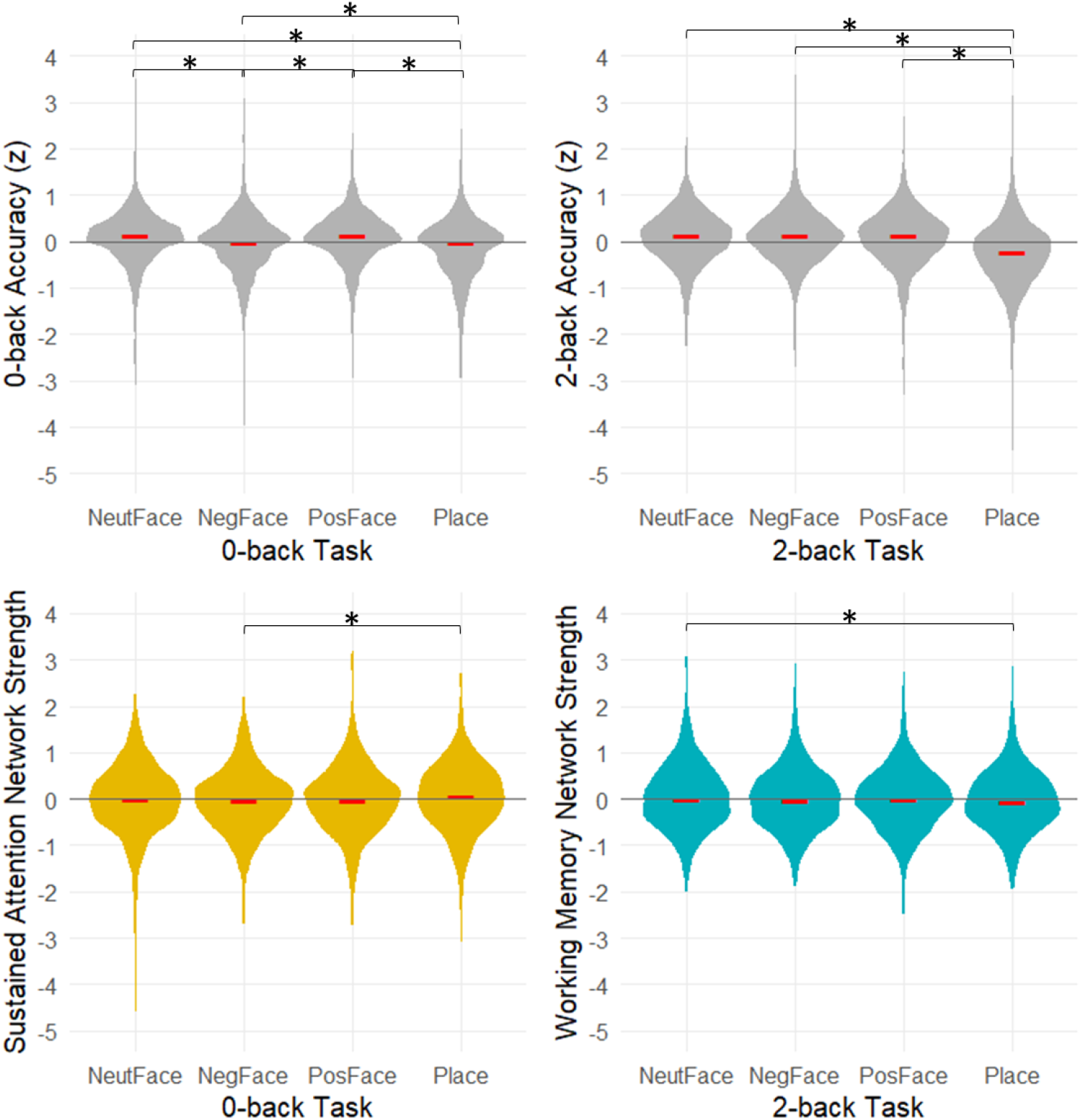
Task accuracy (top) and network strength (bottom) as a function of stimulus types. Red dash indicates median, and * indicates *p*_adj_ < .05 for pairwise comparisons.

**Figure S2.**
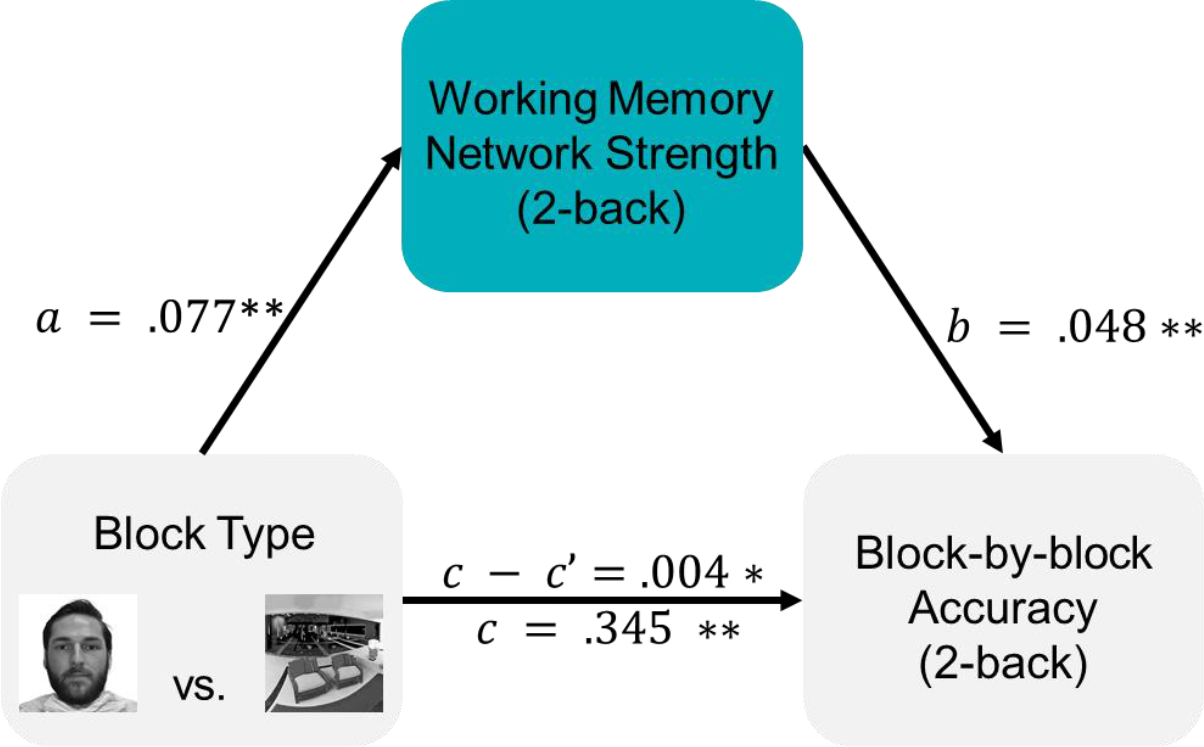
Results of the mediation analyses relating block-by-block performance in the 2-back task to stimulus type, with fluctuations in working memory network strength as a mediator. The total vs. mediated effects are shown as c and c - c’, respectively. All coefficients are adjusted for block-wise motion.

**Figure S3.**
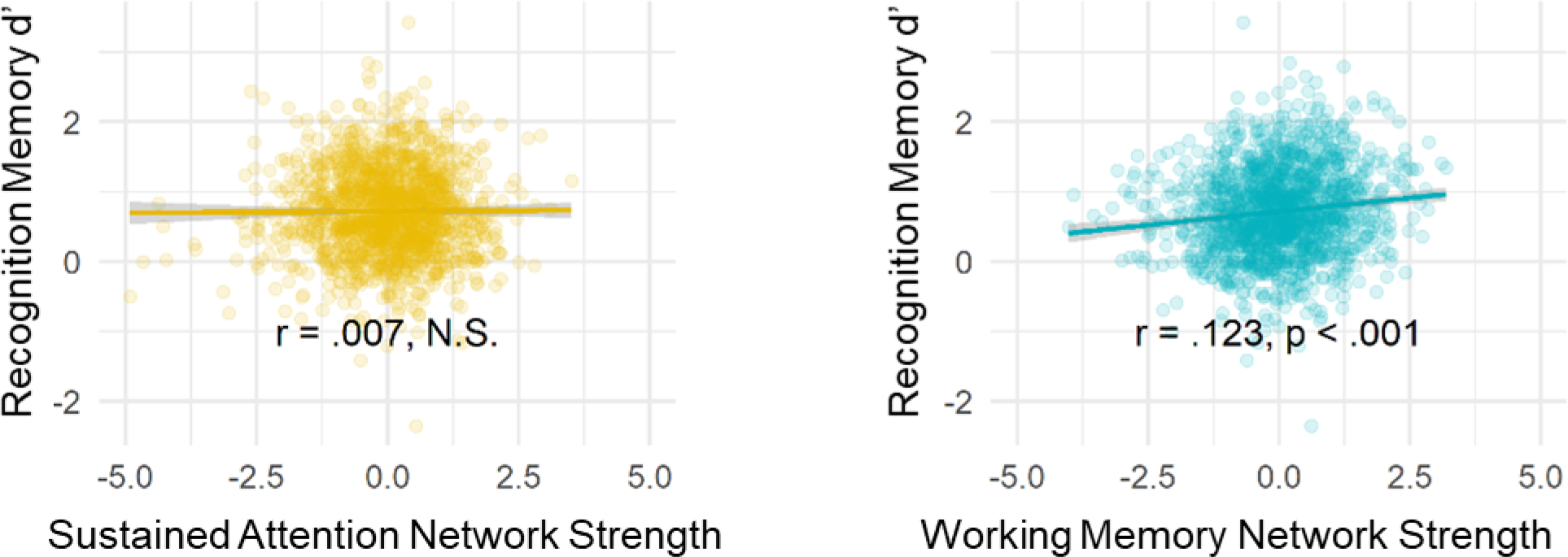
Working memory network strength, but not sustained attention network strength, was predicted children’s subsequent memory task performance.

**Figure S4.**
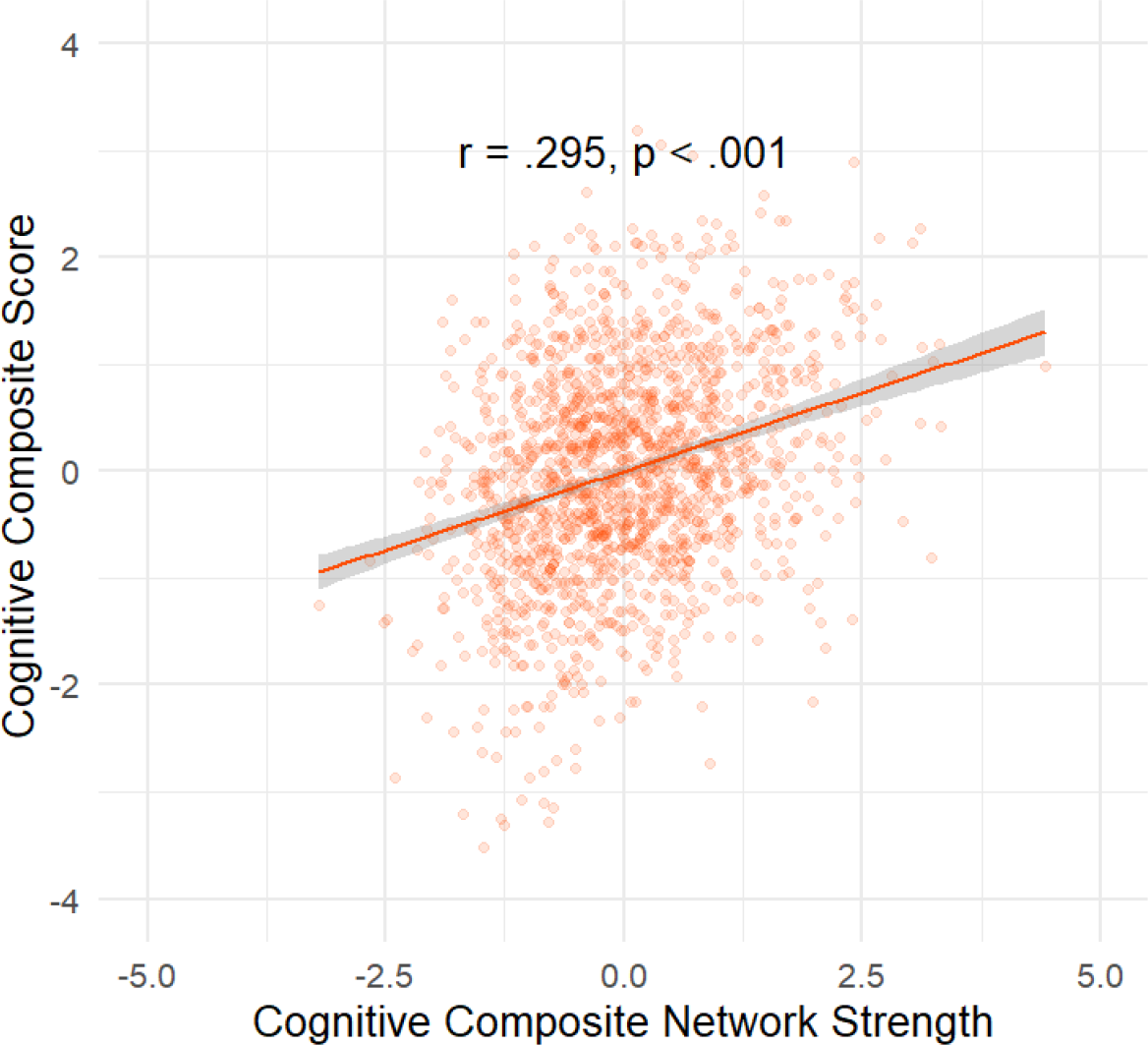
Relationship between cognitive composite network strength and cognitive composite scores in the ABCD sample in all iterations of leave-one-site out combined.

**Figure S5.**
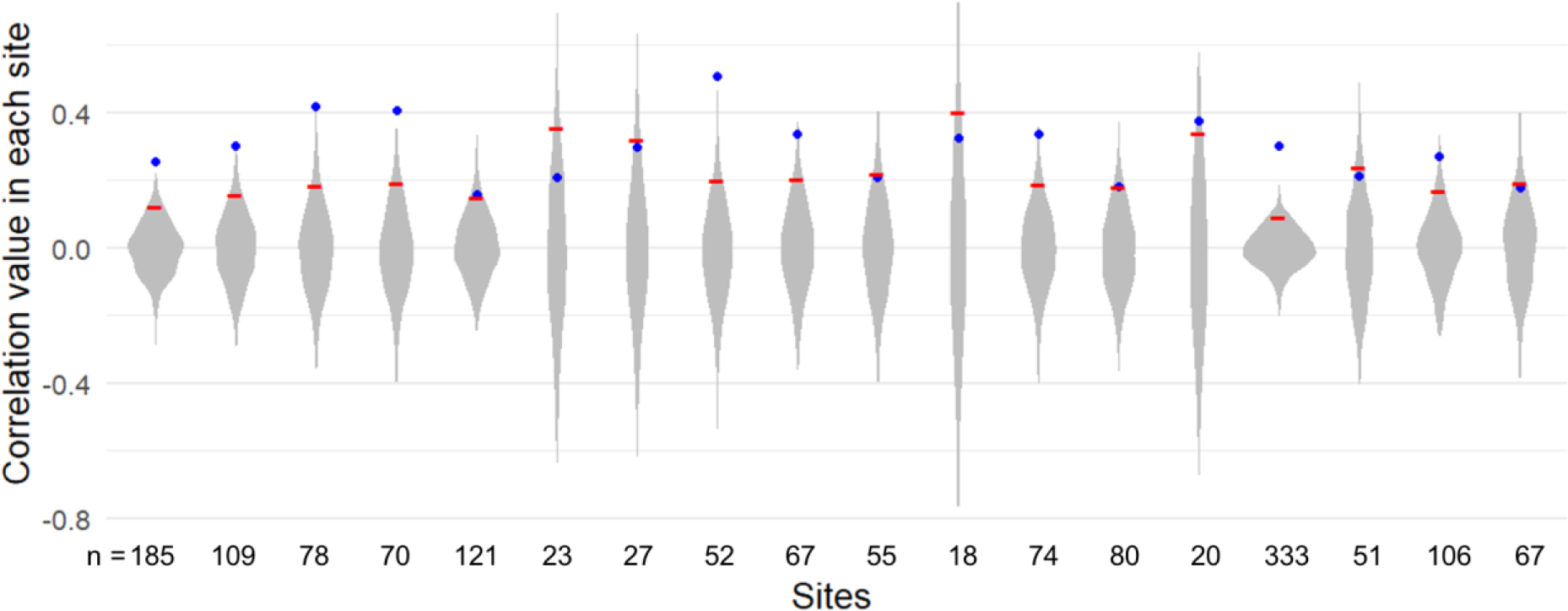
Relationship between cognitive composite network strength and cognitive composite scores in the ABCD sample in each site separately is shown with blue points on permutation null distributions (violins). The red dash indicates the 95^th^ percentile of the null distribution values and the number below each violin indicates the number of participants retained in the study (n) from the corresponding site. Two sites with Philips scanner data and one site with *n* = 3 after motion exclusions were not included in the study analyses (see *Methods*).

**Table S1.** Model predicting individual differences in 0-back and 2-back task accuracy from cognitive composite network strength in the ABCD sample.

| *Predictors* | **0-back Accuracy** |  |  |  | **2-back Accuracy** |  |  |  |
| --- | --- | --- | --- | --- | --- | --- | --- | --- |
|  | *Beta* | *CI* | *t* | *p* | *Beta* | *CI* | *t* | *p* |
| *(Intercept)* | 0.02 | [-0.07, 0.11] | 0.38 | 0.707 | 0.12 | [0.03, 0.21] | 2.67 | **0.012** |
| *Age* | 0.14 | [0.09, 0.19] | 5.75 | **<0.001** | 0.13 | [0.08, 0.17] | 5.22 | **<0.001** |
| *Female* | -0.04 | [-0.18, 0.02] | -0.89 | 0.374 | -0.23 | [-0.32, -0.13] | -4.75 | **<0.001** |
| *Motion* | -0.05 | [-0.10, 0.00] | -1.93 | 0.054 | -0.12 | [-0.16, -0.07] | -4.83 | **<0.001** |
| Cognitive Composite | 0.19 | [0.14, 0.24] | 7.57 | **<0.001** | 0.27 | [0.22, 0.31] | 10.86 | **<0.001** |
| **Random Effects (Site)** |  |  |  |  |  |  |  |  |
| ICC | 0.01<sub>site</sub> |  |  |  | 0.01<sub>site</sub> |  |  |  |
| Observations | 1545 |  |  |  | 1545 |  |  |  |
| Marginal R<sup>2</sup> /<br>Conditional R<sup>2</sup> | 0.071 / 0.083 |  |  |  | 0.136 / 0.148 |  |  |  |

**Table S2.**
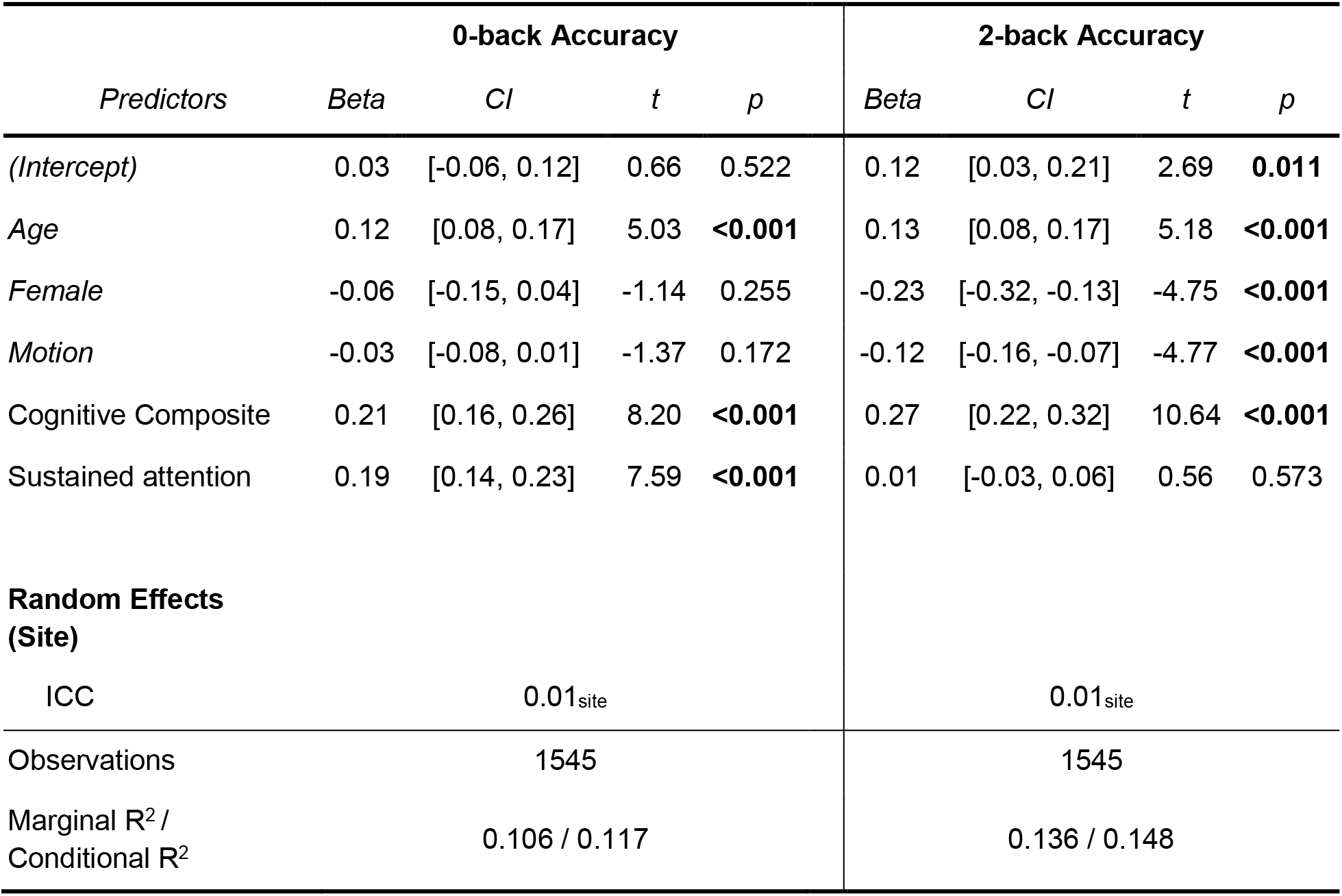
Model predicting individual differences in 0-back and 2-back task accuracy from cognitive composite and sustained attention network strength in the ABCD sample.

| *Predictors* | **0-back Accuracy** |  |  |  | **2-back Accuracy** |  |  |  |
| --- | --- | --- | --- | --- | --- | --- | --- | --- |
|  | *Beta* | *CI* | *t* | *p* | *Beta* | *CI* | *t* | *p* |
| *(Intercept)* | 0.03 | [-0.06, 0.12] | 0.66 | 0.522 | 0.12 | [0.03, 0.21] | 2.69 | **0.011** |
| *Age* | 0.12 | [0.08, 0.17] | 5.03 | **<0.001** | 0.13 | [0.08, 0.17] | 5.18 | **<0.001** |
| *Female* | -0.06 | [-0.15, 0.04] | -1.14 | 0.255 | -0.23 | [-0.32, -0.13] | -4.75 | **<0.001** |
| *Motion* | -0.03 | [-0.08, 0.01] | -1.37 | 0.172 | -0.12 | [-0.16, -0.07] | -4.77 | **<0.001** |
| Cognitive Composite | 0.21 | [0.16, 0.26] | 8.20 | **<0.001** | 0.27 | [0.22, 0.32] | 10.64 | **<0.001** |
| Sustained attention | 0.19 | [0.14, 0.23] | 7.59 | **<0.001** | 0.01 | [-0.03, 0.06] | 0.56 | 0.573 |
| **Random Effects**<br>**(Site)** |  |  |  |  |  |  |  |  |
| ICC | 0.01<sub>site</sub> |  |  |  | 0.01<sub>site</sub> |  |  |  |
| Observations | 1545 |  |  |  | 1545 |  |  |  |
| Marginal R<sup>2</sup> /<br>Conditional R<sup>2</sup> | 0.106 / 0.117 |  |  |  | 0.136 / 0.148 |  |  |  |

**Table S3.** Model predicting intra-individual differences in 0-back and 2-back task accuracy from youth cognitive composite network strength in the ABCD sample.

| *Predictors* | **0-back Accuracy** |  |  |  | **2-back Accuracy** |  |  |  |
| --- | --- | --- | --- | --- | --- | --- | --- | --- |
|  | *Beta* | *CI* | *t* | *p* | *Beta* | *CI* | *t* | *p* |
| *(Intercept)* | 0.07 | [0.02, 0.12] | 2.83 | **0.005** | 0.03 | [-0.02, 0.08] | 1.23 | 0.219 |
| *Block Motion* | -0.03 | [-0.06, -0.01] | -3.20 | **0.001** | -0.02 | [-0.04, -0.00] | -2.11 | **0.035** |
| *Block Type:* |  |  |  |  |  |  |  |  |
| *Negative Face* | -0.13 | [-0.18, -0.09] | -5.80 | **<0.001** | -0.00 | [-0.05, 0.04] | -0.13 | 0.896 |
| *Positive Face* | 0.01 | [-0.03, 0.06] | 0.47 | 0.635 | -0.03 | [-0.07, 0.02] | -1.10 | 0.269 |
| *Place* | -0.23 | [-0.27, -0.18] | -9.99 | **<0.001** | -0.37 | [-0.41, -0.32] | -16.18 | **<0.001** |
| *Run 2 – Run 1* | 0.03 | [-0.01, 0.06] | 1.38 | 0.167 | 0.16 | [0.12, 0.20] | 8.54 | **<0.001** |
| Cognitive Composite | 0.05 | [0.03, 0.07] | 5.37 | **<0.001** | 0.08 | [0.06, 0.10] | 8.33 | **<0.001** |
| **Random Effects (Subject)** |  |  |  |  |  |  |  |  |
| ICC | 0.40<sub>subs</sub> |  |  |  | 0.38<sub>subs</sub> |  |  |  |
| Observations (blocks) | 9176 |  |  |  | 9176 |  |  |  |
| Marginal R<sup>2</sup> /<br>Conditional R<sup>2</sup> | 0.013 / 0.411 |  |  |  | 0.035 / 0.404 |  |  |  |

**Figure S6.**
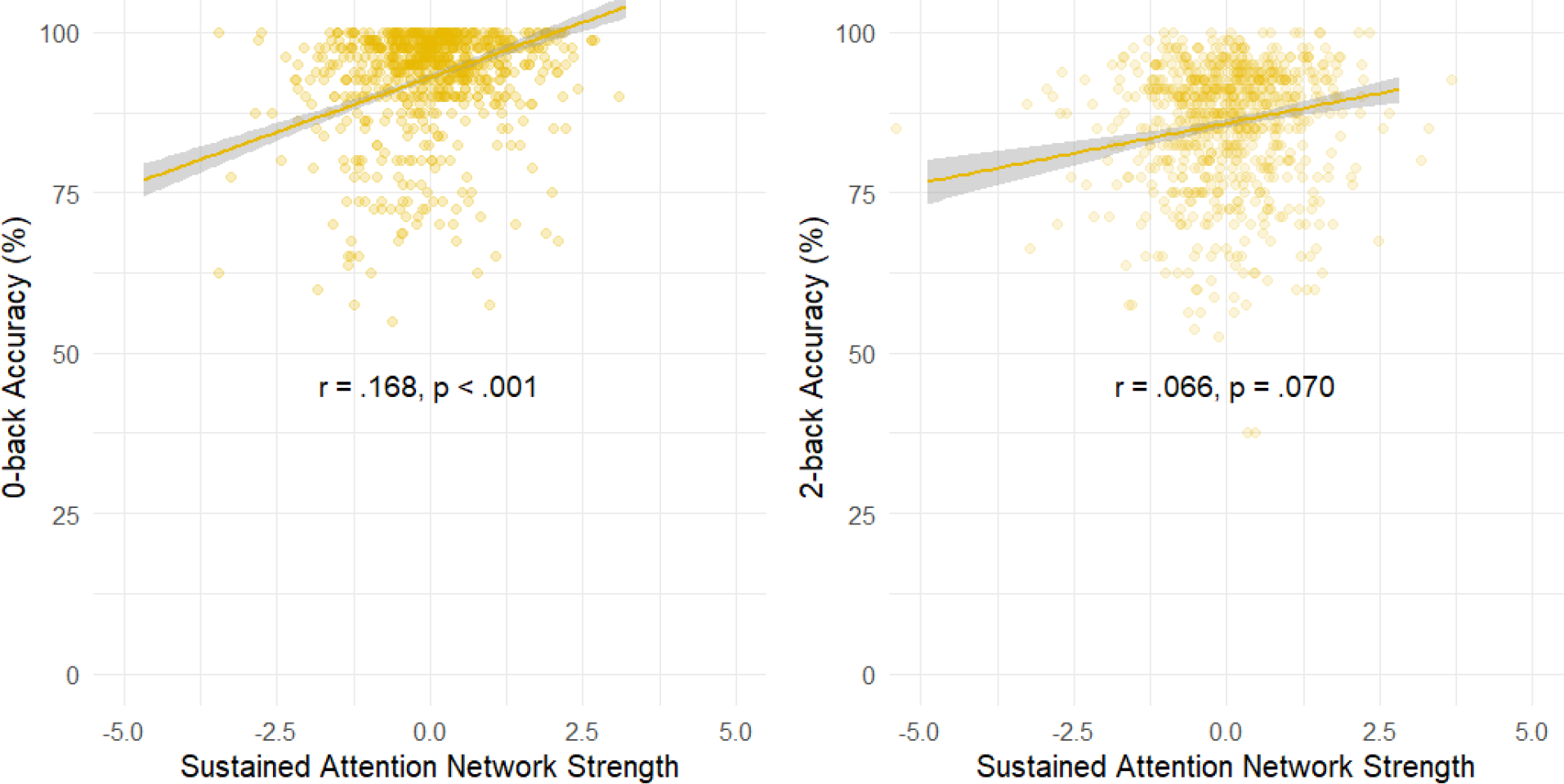
Relationship between sustained attention network strength and *n*-back accuracy in the adult HCP sample.

**Table S4.** Model predicting individual differences in 0-back and 2-back task accuracy from sustained attention network strength in the HCP sample.

| *Predictors* | **0-back Accuracy** |  |  |  |  | **2-back Accuracy** |  |  |  |
| --- | --- | --- | --- | --- | --- | --- | --- | --- | --- |
|  | *Beta* | *CI* | *t* | *p* |  | *Beta* | *CI* | *t* | *p* |
| (Intercept) | 0.07 | [-0.09, 0.22] | 0.86 | 0.390 |  | 0.20 | [0.04, 0.35] | 2.44 | **0.015** |
| Age: |  |  |  |  |  |  |  |  |  |
| Age 26-30 | 0.10 | [-0.09, 0.28] | 1.03 | 0.304 |  | 0.03 | [-0.16, 0.21] | 0.30 | 0.768 |
| Age 31-35 | -0.16 | [-0.35, 0.04] | -1.59 | 0.113 |  | -0.21 | [-0.41, -0.02] | -2.14 | **0.033** |
| Age 36+ | 0.02 | [-0.60, 0.65] | 0.07 | 0.941 |  | -0.31 | [-0.75, 0.50] | -0.40 | 0.691 |
| Female | -0.11 | [-0.25, 0.04] | -1.46 | 0.144 |  | -0.25 | [-0.40, -0.11] | -3.41 | **0.001** |
| Motion | -0.08 | [-0.15, -0.01] | -2.29 | **0.022** |  | -0.09 | [-0.16, -0.02] | -2.51 | **0.012** |
| Sustained Attention | 0.16 | [0.09, 0.23] | 4.51 | **<0.001** |  | 0.07 | [-0.00, 0.14] | -1.95 | 0.051 |
| Observations | 754 |  |  |  |  | 754 |  |  |  |
| Marginal R<sup>2</sup> /<br>Conditional R<sup>2</sup> | 0.053 / 0.046 |  |  |  |  | 0.049 / 0.041 |  |  |  |

**Table S5.** Model predicting intra-individual differences in 0-back and 2-back task accuracy from sustained attention network strength and block type in the HCP sample.

| 0-back Accuracy |  |  |  |  | 2-back Accuracy |  |  |  |
| --- | --- | --- | --- | --- | --- | --- | --- | --- |
| *Predictors* | *Beta* | *CI* | *t* | *p* | *Beta* | *CI* | *t* | *p* |
| *(Intercept)* | 0.12 | [-0.02, 0.25] | 1.73 | 0.083 | -0.01 | [-0.13, 0.12] | -0.11 | 0.912 |
| *Block Motion* | -1.83 | [-2.70, -0.96] | -9.26 | **<0.001** | -1.41 | [-2.20, -0.63] | -3.53 | **<0.001** |
| *Block Type:* |  |  |  |  |  |  |  |  |
| *Face* | 0.38 | [0.32, 0.45] | -12.29 | **<0.001** | 0.68 | [0.62, 0.73] | 23.76 | **<0.001** |
| *Place* | 0.31 | [-0.25, 0.37] | 9.83 | **<0.001** | 0.64 | [0.58, 0.69] | 22.62 | **<0.001** |
| *Tools* | 0.22 | [0.16, 0.28] | 6.98 | **<0.001** | 0.47 | [0.41, 0.52] | -16.34 | **<0.001** |
| *Run 2 – Run 1* | 0.21 | [0.17, 0.26] | 9.26 | **<0.001** | 0.49 | [0.45, 0.53] | 23.29 | **<0.001** |
| Sustained Attention | 0.08 | [0.06, 0.11] | 6.36 | **<0.001** | 0.01 | [-0.02, 0.03] | 0.59 | 0.558 |
| **Random Effects**<br>**(Subject)** |  |  |  |  |  |  |  |  |
| ICC |  | 0.31<sub>subs</sub> |  |  |  | 0.37<sub>subs</sub> |  |  |
| Observations<br>(blocks) |  | 5412 |  |  |  | 5412 |  |  |
| Marginal R<sup>2</sup>/<br>Conditional R<sup>2</sup> |  | 0.046 / 0.344 |  |  |  | 0.135 / 0.456 |  |  |

**Figure S7.**
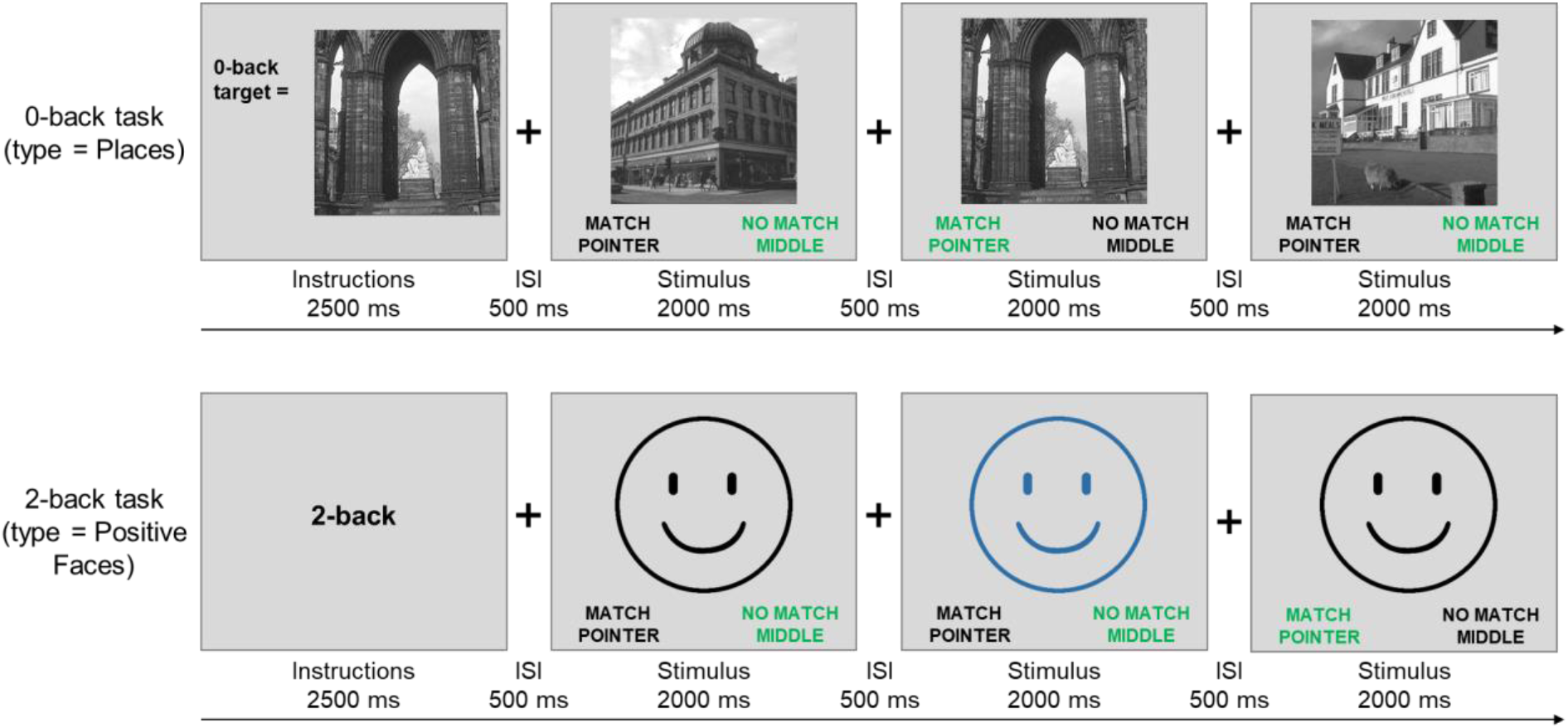
The instruction and first 3 trials in a block of 0-back task (top, example from Places block type) and 2-back task (bottom, example from Positive Faces block type) are shown in this figure. The correct choice for each trial is indicated in green text. Figure adapted from Casey et al., 2018. The real face images are replaced by icons as per *bioRxiv* requirements.

**Table S6.** Model predicting inter-individual differences in 0-back and 2-back task accuracy from sustained attention and working memory network strength values in the ABCD sample with only one of each pair of family members retained randomly (*n* = 1504). Compare these values to Table 1 in the *Results* section.

| 0-back Accuracy |  |  |  |  | 2-back Accuracy |  |  |  |
| --- | --- | --- | --- | --- | --- | --- | --- | --- |
| *Predictors* | *Beta* | *CI* | *t* | *p* | *Beta* | *CI* | *t* | *p* |
| *(Intercept)* | -0.00 | [-0.14, 0.13] | -0.03 | 0.978 | 0.10 | [-0.01, 0.21] | 1.76 | 0.091 |
| *Age* | 0.13 | [0.08, 0.18] | 5.23 | **<0.001** | 0.14 | [0.09, 0.19] | 5.55 | **<0.001** |
| *Female* | -0.06 | [-0.16, 0.04] | -1.19 | 0.236 | -0.22 | [-0.32, -0.12] | -4.38 | **<0.001** |
| *Motion* | -0.03 | [-0.08, 0.02] | -1.03 | 0.304 | -0.13 | [-0.18, -0.08] | -5.14 | **<0.001** |
| Sustained Attention | 0.15 | [0.10, 0.20] | 6.10 | **<0.001** | -0.04 | [-0.09, 0.01] | -1.72 | 0.086 |
| Working Memory | 0.11 | [0.06, 0.16] | 4.13 | **<0.001** | 0.10 | [0.05, 0.15] | 3.85 | **<0.001** |
| **Random Effects**<br>**(Site)** |  |  |  |  |  |  |  |  |
| ICC |  | 0.06<sub>site</sub> |  |  |  | 0.03<sub>site</sub> |  |  |
| Observations |  | 1504 |  |  |  | 1545 |  |  |
| Marginal R<sup>2</sup>/<br>Conditional R<sup>2</sup> |  | 0.067 / 0.119 |  |  |  | 0.070 / 0.099 |  |  |

**Table S7.**
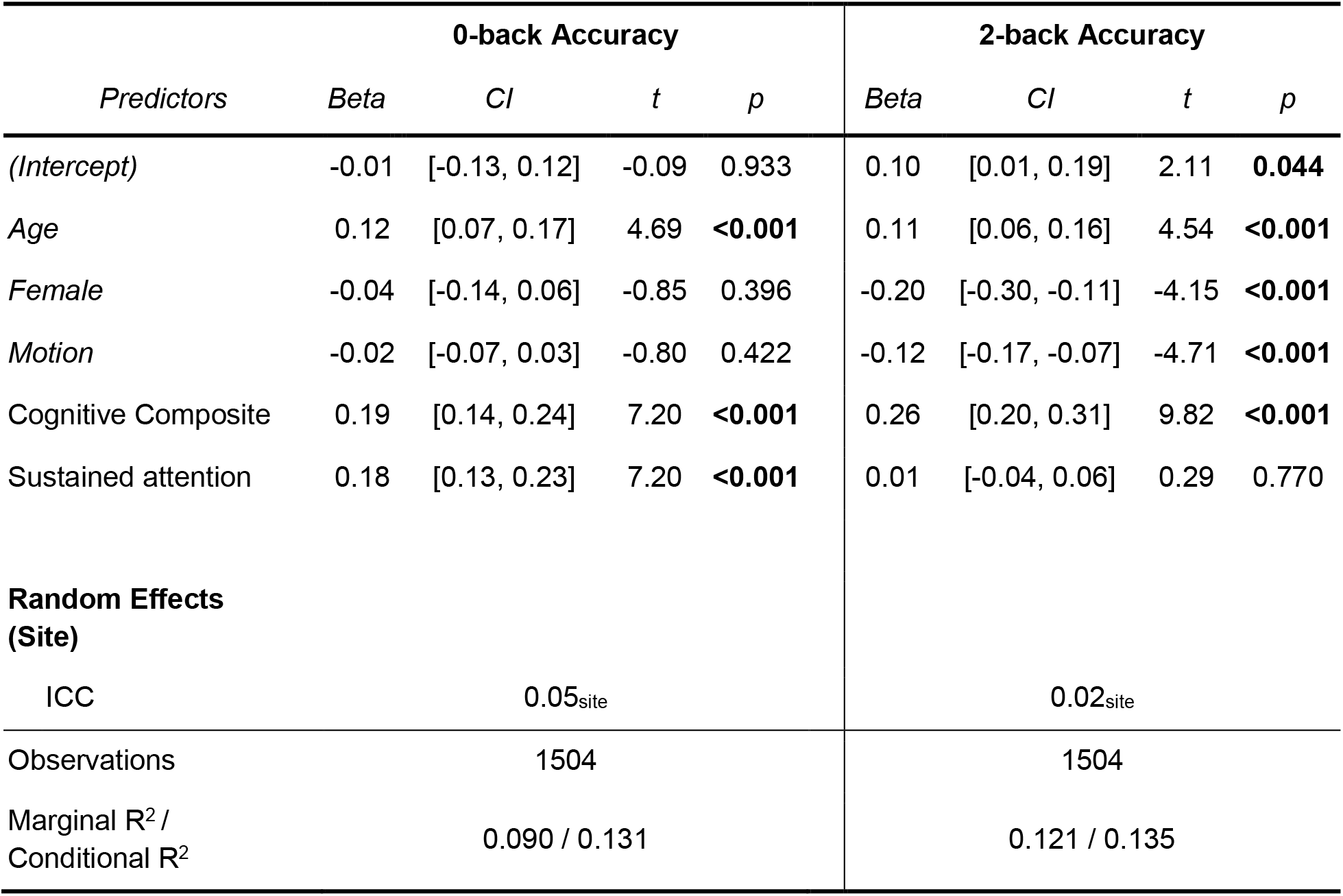
Model predicting inter-individual differences in 0-back and 2-back task accuracy from sustained attention and youth cognitive composite network strengths in the ABCD sample with only one of each pair of family members retained randomly (*n* = 1504). Compare these to Table S4.

**Table S8.**
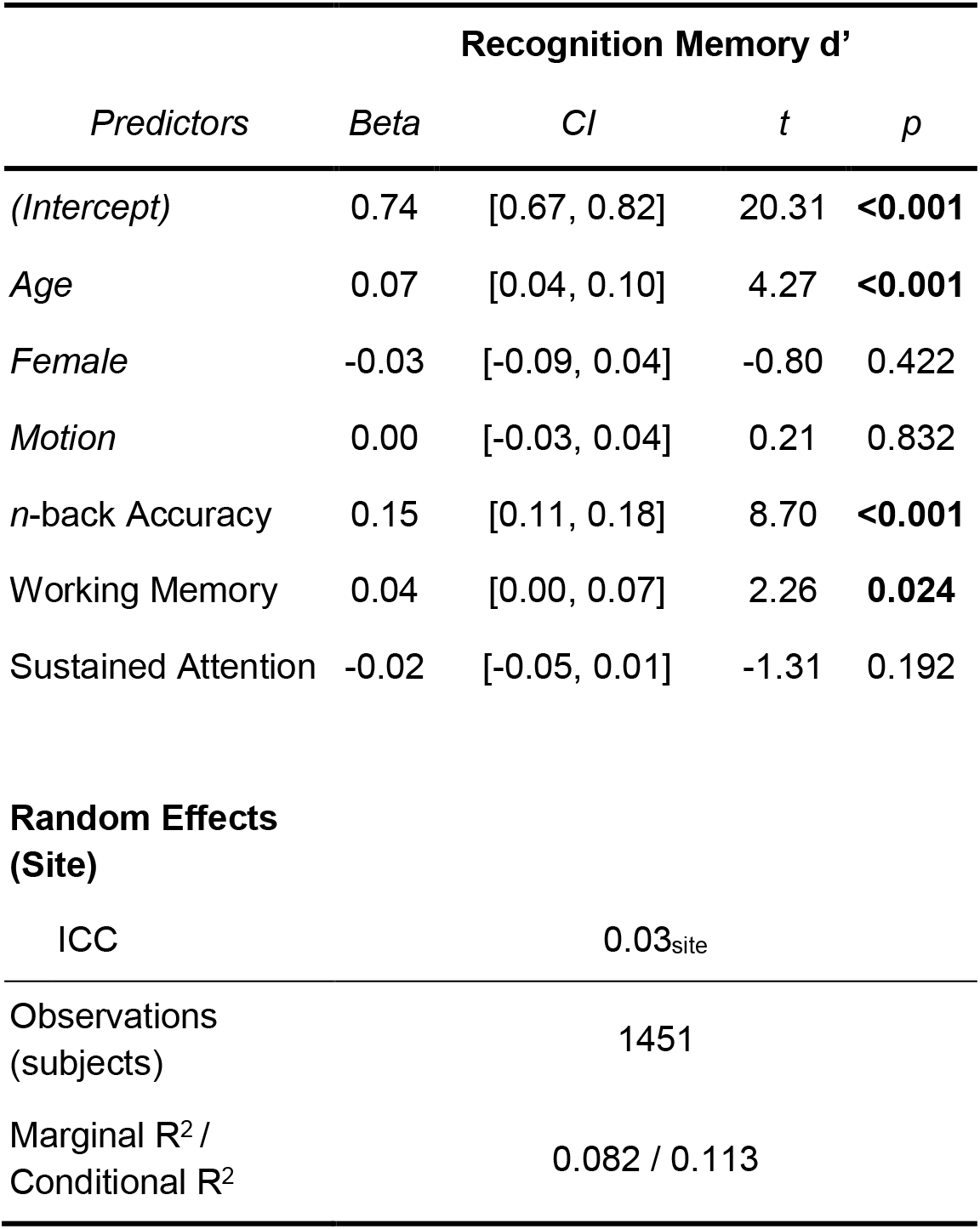
Model predicting out-of-scanner item memory recognition from sustained attention and working memory network strengths in the ABCD sample with only one of each pair of family members retained randomly. Compare these to Table 3.

| Recognition Memory $d'$ | | | | |
| --- | --- | --- | --- | --- |
| *Predictors* | *Beta* | *CI* | *t* | *p* |
| (Intercept) | 0.74 | [0.67, 0.82] | 20.31 | **<0.001** |
| Age | 0.07 | [0.04, 0.10] | 4.27 | **<0.001** |
| Female | -0.03 | [-0.09, 0.04] | -0.80 | 0.422 |
| Motion | 0.00 | [-0.03, 0.04] | 0.21 | 0.832 |
| *n*-back Accuracy | 0.15 | [0.11, 0.18] | 8.70 | **<0.001** |
| Working Memory | 0.04 | [0.00, 0.07] | 2.26 | **0.024** |
| Sustained Attention | -0.02 | [-0.05, 0.01] | -1.31 | 0.192 |

|  |  |
| --- | --- |
| ICC | 0.03<sub>site</sub> |
| Observations<br>(subjects) | 1451 |
| Marginal $R^2$ /<br>Conditional $R^2$ | 0.082 / 0.113 |

### Supplementary analysis: Defining *n*-back blocks with a temporal lag

In each of the two *n*-back runs, there were 8 blocks of 0- and 2-back tasks (4 of each, one per stimulus category). All of the main analyses shown in the *Results* use block-wise functional connectivity matrices for which block start and end times were the first stimulus onset and last stimulus offset times for trials in the block, respectively. However, the rest period between blocks were not equally distributed for all 8 blocks in a run, and instead followed the design shown in the Figure S8. As shown in the figure, there was enough time before the odd blocks (i.e., blocks 1, 3, 5, and 7) for the BOLD response from the previous *n*-back block to return to baseline, but this was not the case for the even blocks (i.e., blocks 2, 4, 6, 8). As such, in a post hoc analysis we calculated functional connectivity matrices for the even blocks with a 6-s shift (7 or 8 TRs based on rounding the exact timings for each participant) to allow for further separation of the BOLD signal from these blocks with their prior blocks. This is shown in Figure S8 red intervals (compared to blue intervals in the analyses presented in the *Results* sections). We then repeated all ABCD analyses using these shifted functional connectivity matrices. Results were similar to those reported in the non-shifted analysis, except for the **Study 1.3** results shown in Table 3. Therefore, we report those results with the time-shifted functional connectivity matrices here. With the time-shifted matrices, working memory network strength still predicted subsequent recognition memory for *n*-back stimuli, although the strength of the association was weaker (*r* = .093, *p* < .001 vs. *r* = .123 in original analysis) and was no longer significant above and beyond in-scanner *n*-back performance (shown in Table S8).

**Figure S8.**
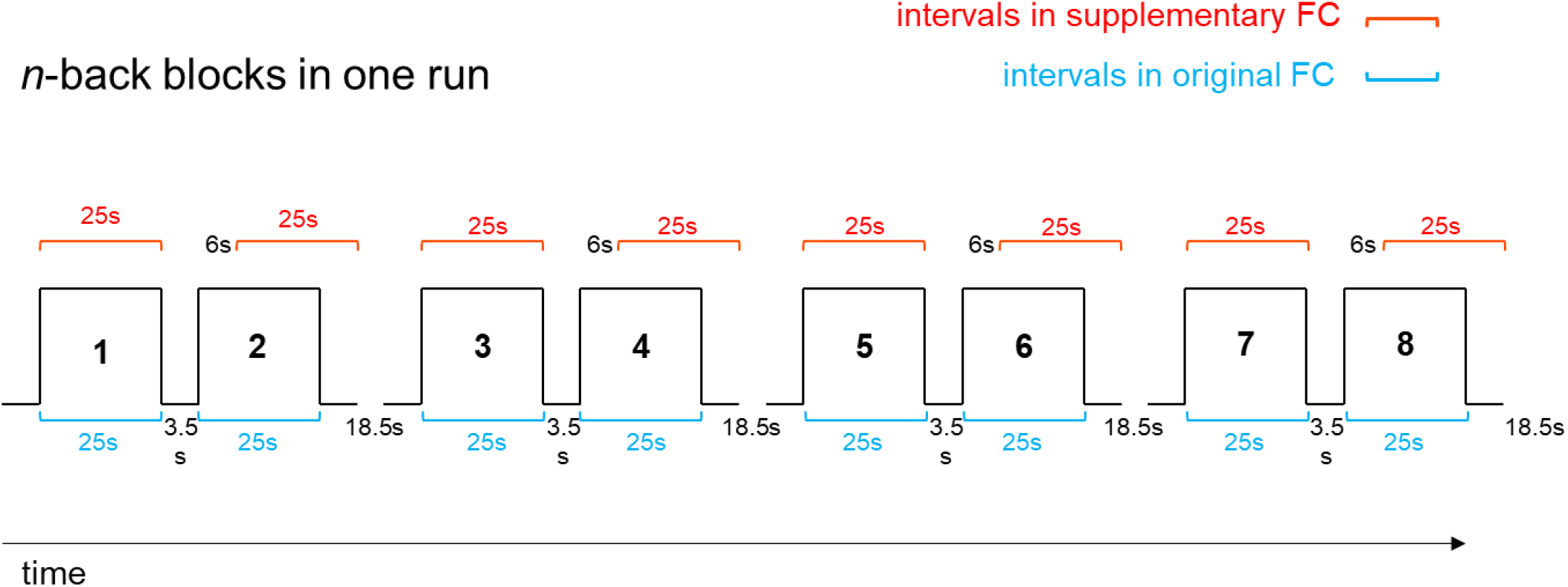
The time-course of *n*-back blocks (from the beginning of the first trial’s stimulus onset to the end of the last trial’s stimulus offset time in the block) in a run in the ABCD n-back task. While there is only ∼3.5-sec interval prior to start of even blocks, there is an additional 15-sec rest interval after every other block of *n*-back.

**Table S9.** Working memory network strength measured in the 6-sec shifted manner described above is not significantly related to subsequent recognition memory for *n*-back task stimuli after adjusting for nuisance variables and *n*-back performance itself (compare to Table 3).

| Recognition Memory $d'$ | | | | |
| --- | --- | --- | --- | --- |
| *Predictors* | *Beta* | *CI* | *t* | *p* |
| *(Intercept)* | 0.73 | [0.66, 0.79] | 23.22 | **<0.001** |
| *Age* | 0.06 | [0.03, 0.09] | 3.97 | **<0.001** |
| *Female* | -0.02 | [-0.08, 0.04] | -0.66 | 0.511 |
| *Motion* | 0.01 | [-0.02, 0.04] | 0.73 | 0.468 |
| *n*-back Accuracy | 0.18 | [0.15, 0.21] | 10.85 | **<0.001** |
| Working Memory | 0.03 | [-0.01, 0.06] | 1.59 | 0.113 |
| Sustained Attention | -0.02 | [-0.05, 0.01] | -1.34 | 0.179 |
| **Random Effects** |  |  |  |  |
| **(Site)** |  |  |  |  |
| ICC | 0.02<sub>site</sub> |  |  |  |
| Observations<br>(subjects) | 1489 |  |  |  |
| Marginal $R^2$ /<br>Conditional $R^2$ | 0.102 / 0.118 | | | |

## Footnotes

1 The correlation of recognition memory *d’* with 0-back and 2-back accuracy separately is *r* = .26 and *r* = .30, respectively.

2 Cross-block covariance σ_XY_ is estimated as the ratio of the LV’s squared eigenvalue over the sum of squared eigenvalues across all the LVs and represents the dominance of an LV in the PLS analysis.

